# Conserved and Divergent Modulation of Calcification in Atherosclerosis and Aortic Valve Disease by Tissue Extracellular Vesicles

**DOI:** 10.1101/2020.04.02.022525

**Authors:** Mark C. Blaser, Fabrizio Buffolo, Arda Halu, Florian Schlotter, Hideyuki Higashi, Lorena Pantano, Louis A. Saddic, Samantha K. Atkins, Maximillian A. Rogers, Tan Pham, Eugenia Shvartz, Galina K Sukhova, Silvia Monticone, Giovanni Camussi, Simon C. Body, Jochen D. Muehlschlegel, Sasha A. Singh, Masanori Aikawa, Elena Aikawa

## Abstract

**Background:** Fewer than 50% of patients develop calcification of both atherosclerotic plaques and aortic valves, implying differential pathogenesis. While circulating extracellular vesicles (EVs) act as biomarkers of cardiovascular diseases, tissue-entrapped EVs associate with early mineralization, but their contents, function, and contributions to disease remain unknown.

**Results:** Global proteomics of human carotid artery endarterectomies and calcified aortic valves from a total of 27 donors/patients revealed significant over-representation of proteins with vesicle-associated pathways/ontologies common to both diseases. We exploited enzymatic digestion, serial (ultra)centrifugation and OptiPrep density-gradient separation to isolate EV populations from diseased arteries and valves. Mass spectrometry found 22 EV marker proteins to be highly enriched in the four least-dense OptiPrep fractions while extracellular matrix proteins predominated in denser fractions, as confirmed by CD63 immunogold electron microscopy and nanoparticle tracking analysis. Proteomics and miRNA-sequencing of OptiPrep-enriched tissue EVs quantified 1,104 proteins and 123 miR cargoes linked to 5,182 target genes. Pathway networks of proteins and miR targets common to artery and valve tissue EVs revealed a shared regulation of Rho GTPase and MAPK intracellular signaling cascades. 179 proteins and 5 miRs were significantly altered between artery and valve EVs; multi-omics integration determined that EVs differentially modulated cellular contraction and p53-mediated transcriptional regulation in diseased vascular vs. valvular tissue.

**Conclusions:** Our findings delineate a strategy to isolate, purify, and study protein and RNA cargoes from EVs entrapped in fibrocalcific tissues. Multi-omics and network approaches implicated tissue-resident EVs in human cardiovascular disease.

## Background

Cardiovascular calcification directly correlates with incidence of cardiovascular events such as myocardial infarction, stroke, and heart failure, and strongly predicts morbidity and mortality.(1) While aberrant mineralization of arteries and heart valves share numerous risk factors, only 25-50% of patients develop both vascular and valvular calcification, implying that they involve differential drivers.(2) Histopathological studies described these two conditions as being grossly comparable (3) and there has been little subsequent study of this apparent epidemiological paradox. Despite the histological similarities, pharmacological therapies such as statins have been successful in mitigating inflammation and lowering cholesterol in patients with atherosclerosis but they failed to improve outcomes for aortic valve stenosis – leaving patients without effective therapeutics and lending further credence to the notion that pathological changes, including calcification, follow diverse routes in these tissue beds.(4) Calcification-prone extracellular vesicles (EVs) offer a possible explanation: recent work has implicated EVs as important drivers and building blocks of mineralization in the vasculature.(5–7, 8) EVs are small lipid membrane-bound structures secreted by all human cell types.(9) Found throughout the body’s tissues and fluids, these particles arise from two regulated paths: exosomes (~50-150 nm in diameter) are secreted when multivesicular bodies of endosomal origin fuse with the plasma membrane, and microvesicles (~100-500 nm in diameter) are produced by budding and fission of the cell membrane (reviewed in (10)). EVs contain actively-selected bioactive cargoes (lipids, proteins, and noncoding microRNAs [miRs]) and mediate cell-cell communication.(11) Circulating plasma levels of EVs rise in patients with coronary artery disease and associate with a higher risk of cardiovascular death, revascularization, and major adverse cardiovascular and cerebral event occurrence.(12) These findings have spurred a focus on circulating EV cargoes as biomarkers in cardiovascular disease.

Although EVs may arise from endothelial cells or platelets, cell culture experiments have revealed that EVs from other cell types found in diseased vessels and/or valves may directly drive calcification.(13) Cultured macrophages release EVs with high aggregation potential that nucleate hydroxyapatite, and EVs derived from SMCs grown under calcifying conditions are loaded with pro-mineralizing annexins and phosphatidylserine along with reduced levels of calcification inhibitors.(6, 14) As a result, they act as sites of mineral nucleation and modulate the formation of early plaque microcalcifications.(8) Recent work has localized EVs in both human atherosclerotic plaques (6, 8) and calcified vessels from patients with chronic kidney disease.(15, 16) As in other microenvironments, such tissue-entrapped EVs presumably mediate intercellular signaling within the plaque, but they have not been previously characterized. Unlike the routine isolation of EVs from biofluids, extracting tissue EVs that are of sufficient purity, quality, and quantity for downstream applications presents substantial challenges.(17) As such, tissue EV contents have only been comprehensively studied in more readily disrupted human brain and murine lung tumors.(17, 18) Tissue-entrapped EVs therefore are a rich, untapped source of biological insight into putative mechanisms and therapeutic targets of diseases.

The present study hypothesized that tissue-entrapped vesicular cargoes contribute to the shared- and tissue-specific pathogenesis of cardiovascular calcification. Using global proteomics, we examined EVs in whole-tissue samples from calcified human carotid artery plaques and aortic valves. Guided by mass spectrometric measures of separation performance, we then developed a hybrid isolation approach that employed enzymatic digestion to release extracellular matrix (ECM)/tissue-bound EVs, followed by sequential multi-step centrifugation, ultracentrifugation, and OptiPrep density gradient separation to remove non-EV contaminants and enrich the EV population (Figure 1). Lastly, we applied proteomics, small RNA sequencing, multi-omics integration, multi-dimensional network analysis, and bioinformatic techniques to derive biological insights from tissue-entrapped vesicle cargoes to assess EVs as regulators of the differential pathogenesis of vascular and valvular calcification.

**Figure 1:**
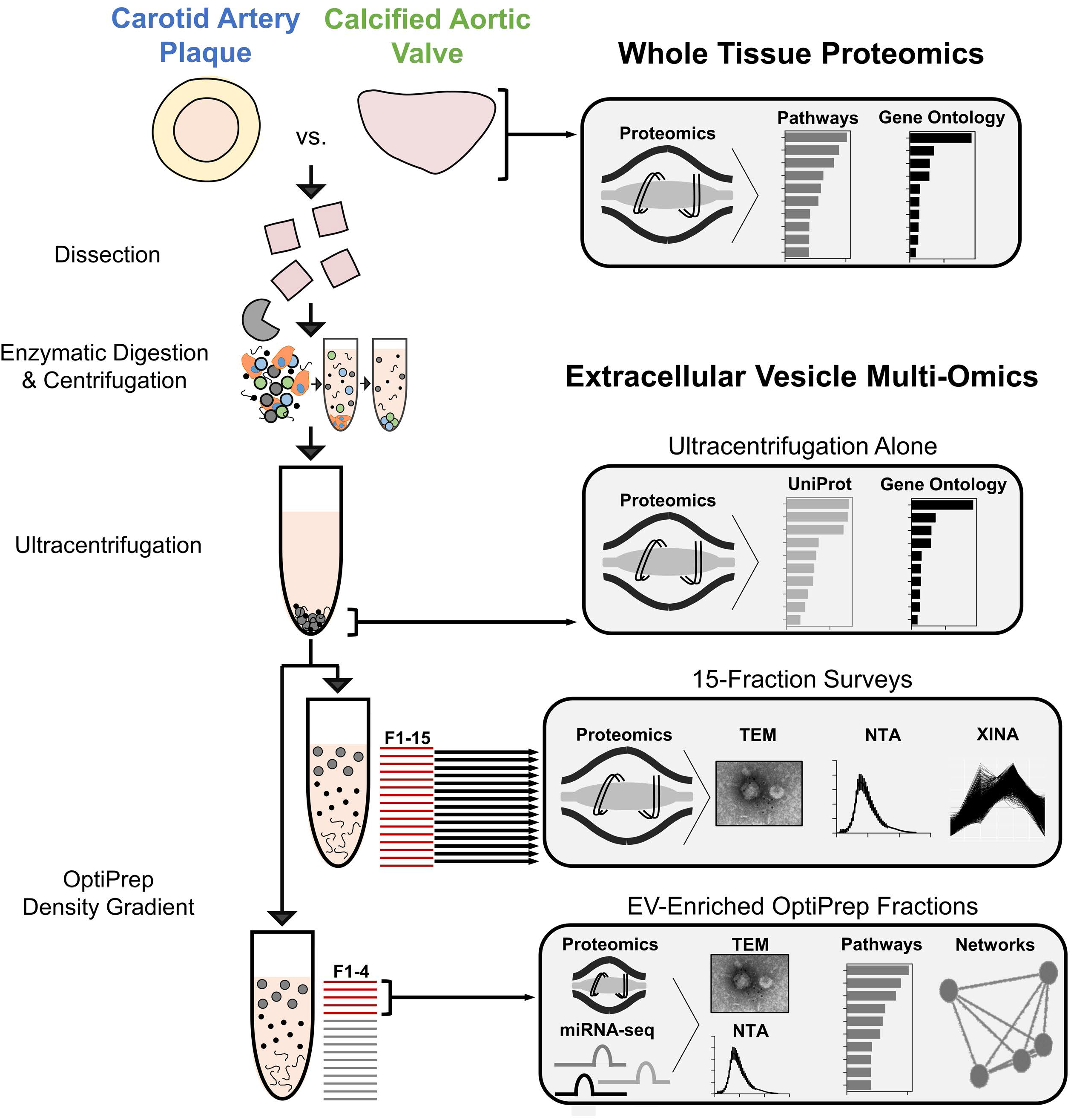
Experimental Overview – Isolation and Analysis of Cardiovascular Tissue-Entrapped Extracellular Vesicles. Label-free proteomics was conducted on *whole tissue* samples of fresh human carotid artery plaques or calcified aortic valves. Alternatively, samples underwent enzymatic digestion and serial low- and high-speed centrifugation. The high-speed supernatant then underwent ultracentrifugation to wash/pellet EVs. *15-fraction survey* experiments utilized OptiPrep density gradient separation in concert with mass spectrometry, transmission electron microscopy (TEM), nanoparticle tracking analysis (NTA), and coabundance profiling (XINA) to identify which fractions were enriched in EVs and which contained non-EV contaminants. Enrichment performance was compared to that of *ultracentrifugation alone* by performing mass spectrometry on the pellet resulting from ultracentrifuged tissue digests. Tissue EV cargoes were then rigorously assessed by pooling *EV-enriched OptiPrep fractions* (F1-4) together and employing mass spectrometry, transcriptomics, TEM, NTA, and bioinformatics approaches to interrogate and classify these vesicles.

## Results

### Whole-Tissue Proteomics Identifies Conservation of EV-Associated Protein Expression between Advanced Vascular and Valvular Disease

While proteomics has been previously performed separately on atherosclerotic plaques (19, 20) or calcified stenotic aortic valves,(21) the proteomes of these diseased cardiovascular tissues have never been directly compared in any species. To identify shared or deviating mechanisms underlying disease pathogenesis in these two tissues, we performed label-free liquid-chromatography-mass spectrometry (LC-MS) on cryogenically-homogenized human atherosclerotic plaques from carotid endarterectomies and calcified aortic valves that had undergone surgical replacement. Though there was near-complete overlap amongst the 1,678 proteins that we identified by whole tissue proteomics of carotid artery plaques and calcified aortic valves (Figure 2A), 237 proteins were differentially enriched between tissue types (137 proteins up in carotid artery, 130 up in aortic valve; false discovery rate (FDR)≤0.05; Figure 2B) and principal component analyses found clear tissue-specific clustering across the complete proteome. Together, these data indicated the presence of heterogeneity between these forms of advanced cardiovascular disease. Inspection of highly-significant and highly-enriched proteins in carotid artery (Figure 2C) revealed a proteome dominated by associations with cell adhesion, migration, and cytoskeletal force generation (TES, PDLIM4, TINAGL1, FBLIM1, ACTA2, ITGA8); SMC plasticity (LMOD1); ECM synthesis, organization, and degradation (COL14A1, MMP12); inflammation, lipid metabolism and macrophage efferocytosis (IL4l1, AOC3, CAMK2G); and calcification (NPNT, ITGB3). We investigated the global impact of those proteins augmented in carotid artery plaque via pathway enrichment analysis: 188 KEGG, BioCarta, and Reactome pathways (Supplemental Table 1) were significantly enriched and included links to SMC contraction, bone mineralization, triglyceride metabolism, and inflammatory cell regulation/migration/differentiation. In contrast, proteins most significantly enriched in calcified valves (Figure 2E) were associated with neural crest differentiation (MIA, GFAP, SEMA3B; likely due more to the developmental origins of the valve/proximal aorta and less-so to divergent disease); osteochondral differentiation (PTN, COMP, CILP, CILP2, PRG4, CHAD); calcification (ALPL, COMP); proteoglycan sulfation or proteolysis (CHST6, HTRA3); extracellular matrix metabolism (TNXB, PODNL1, FBLN1); and elastin fragmentation (FBN1). Globally, 69 pathways (Supplemental Table 2) were significantly enriched in the valvular proteome, including those involved in proteoglycan biosynthesis and metabolism, TGF-β signaling, detoxification of reactive oxygen species, cell/matrix interactions, and the innate immune system. Importantly, when we examined pathway enrichment in the 1,411 proteins whose abundances were unchanged between diseased vascular and valvular tissue (FDR>0.05), there was a statistically significant 5.08-fold increase in the incidence of vesicle-associated pathways vs. the total pathway database (p<0.01; Figure 2D, Supplemental Table 3). Furthermore, half of the gene ontology terms that were most significantly enriched in this unchanged proteome associated with EVs, exosomes, exocytosis, or secretion (Figure 2D, Supplemental Table 4). Together, these observations implied that EV-related functionality was present in both calcified carotid artery plaques and aortic valves, prompting us to isolate and directly interrogate the contents of these tissue-entrapped EVs.

**Figure 2:**
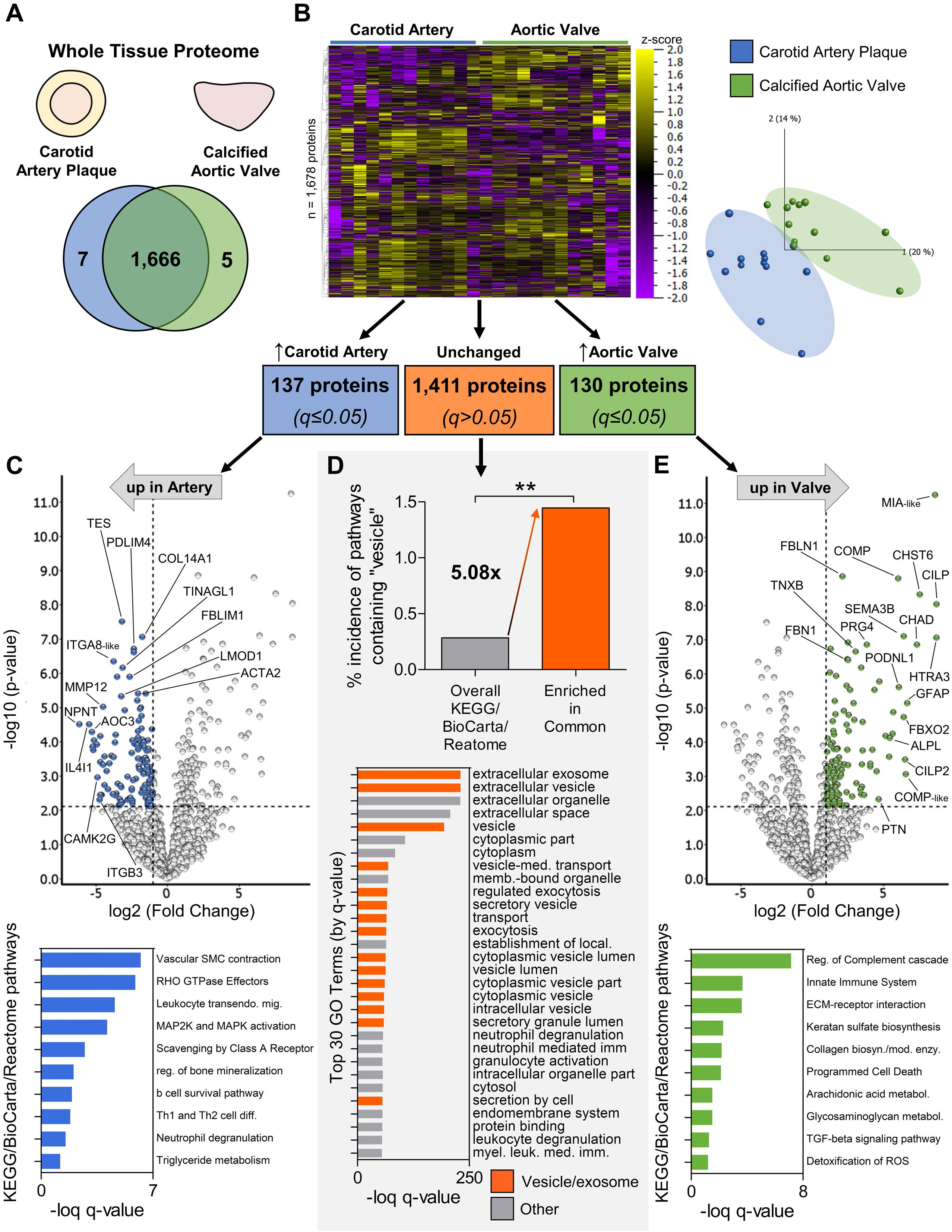
Whole-Tissue Proteomics Finds that Extracellular Vesicle-Associated Functions are Conserved Between Advanced Vascular and Valvular Disease. **A,** The whole-tissue proteome was composed of 1,678 proteins, of which 99.3% were identified in both calcified human carotid artery and aortic valve samples; n=4 carotid artery and 4 aortic valve donors. **B,** Despite this near-complete overlap, principal component analysis demonstrated tissue-type specific clustering of samples and 267 proteins were significantly differentially enriched between carotid artery plaque and calcified aortic valve tissue (proteins filtered at q≤0.05). **C and E,** Top: volcano plots of whole-tissue proteomics data. Blue and green markers indicate proteins with high differential enrichment in carotid artery and aortic valve tissue, respectively, with cutoffs at a fold-change of 2 and a p-value of 0.00795 (corresponding to a q-value of 0.05). Bottom: selected KEGG, Reactome, and BioCarta pathways that were significantly enriched among those proteins significantly increased in calcified carotid artery plaques (C, 137 proteins) and aortic valve tissue (E, 130 proteins). **D,** In pathways enriched in the portion of the whole-tissue proteome whose abundances were unchanged between tissue types (1,411 proteins, q>0.05), we found a found a statistically significant 5.08-fold increase in the incidence of vesicle-associated pathways vs. the total pathway database (p=0.0024). In addition, 15 of the top 30 most-significant gene ontology terms in the unchanged whole-tissue proteome were associated with extracellular vesicles, exosomes, exocytosis, or secretion (orange bars).

### Proteomics, Enzymatic Digestion, Serial (ultra)Centrifugation, and Density Gradient Separation Isolate and Enrich Tissue-Entrapped EVs

We dissociated calcified human carotid artery plaques and aortic valves by bacterial collagenase digestion. Then, to enrich and purify EVs from crude tissue digests, we coupled sequential centrifugation and ultracentrifugation to 15-fraction OptiPrep density gradient separation, an EV enrichment approach that has been used with success in biofluids (Supplemental Figure 1). We employed label-free mass spectrometry to survey each of the 15 density gradient fractions in order to assess where EVs had collected. We quantified 1,491 proteins and determined that 22 EV marker proteins (e.g. annexins, CD63/81, flotillins, HSP70s, TSG101, tetraspanins) (10) were consistently enriched in the four least-dense fractions from both calcified carotid artery plaques and aortic valves, peaking in fraction 3 (Figure 3A,C, Supplemental Figure 2A). Using a clustering-to-networks software developed by our laboratory (XINA (22)), we performed a coabundance analysis on the proteins across the 15 OptiPrep fractions. The coabundance profiles demonstrated further indications of EV enrichment in fractions 1-4: proteins contained in the five XINA clusters whose abundance profiles mimicked those of the 22 EV markers were highly significant for gene ontology terms associated with vesicular processes (Supplemental Figure 2B, Supplemental Tables 5,6). Furthermore, OptiPrep fractionation successfully separated EVs from non-EV contaminants, as abundant ECM proteins were detected in denser fractions (Figure 3A,C). Globular collagens (e.g. Collagen VIA1, VIA2, VIA3) and other ECM components (e.g. Fibrillin 1 and Versican) were enriched in fractions 5-10, and fibrillar collagens (e.g. Collagen IA1, IA2, IIIA1) were found primarily in fractions 11-15. This phenotypic fractionation was also visualized by transmission electron microscopy (TEM): CD63-labelled immunogold beads confirmed CD63+ membrane-bound EVs in the low-density OptiPrep fractions of both vascular and valvular tissues, along with globular collagens in fractions 5-10 and fibrillar collagens in the densest fractions (Figure 3E; fraction-by-fraction CD63-labelled TEM and negative controls are shown in Supplemental Figure 3; negative controls to Figure 3E in Supplemental Figure 4). We then employed nanoparticle tracking analysis (NTA) to confirm the presence and quantify the size of EV particles (Figure 3B,D). In fractions 1-4, NTA identified particles ~200 nm in diameter with a size distribution characteristic of EVs, both of which agree with our previous reports of EVs released *in vitro* from cultured macrophages, vascular SMCs, and valve interstitial cells (VICs) under calcifying conditions.(6, 16, 23) While vesicles were enriched in fractions 1-4 (demonstrated by proteomics, TEM, and NTA), mean particle size remained relatively consistent (~100-250 nm) across all fractions (Figure 3B,D), likely due to the presence of digested collagen fragments in denser fractions.

**Figure 3:**
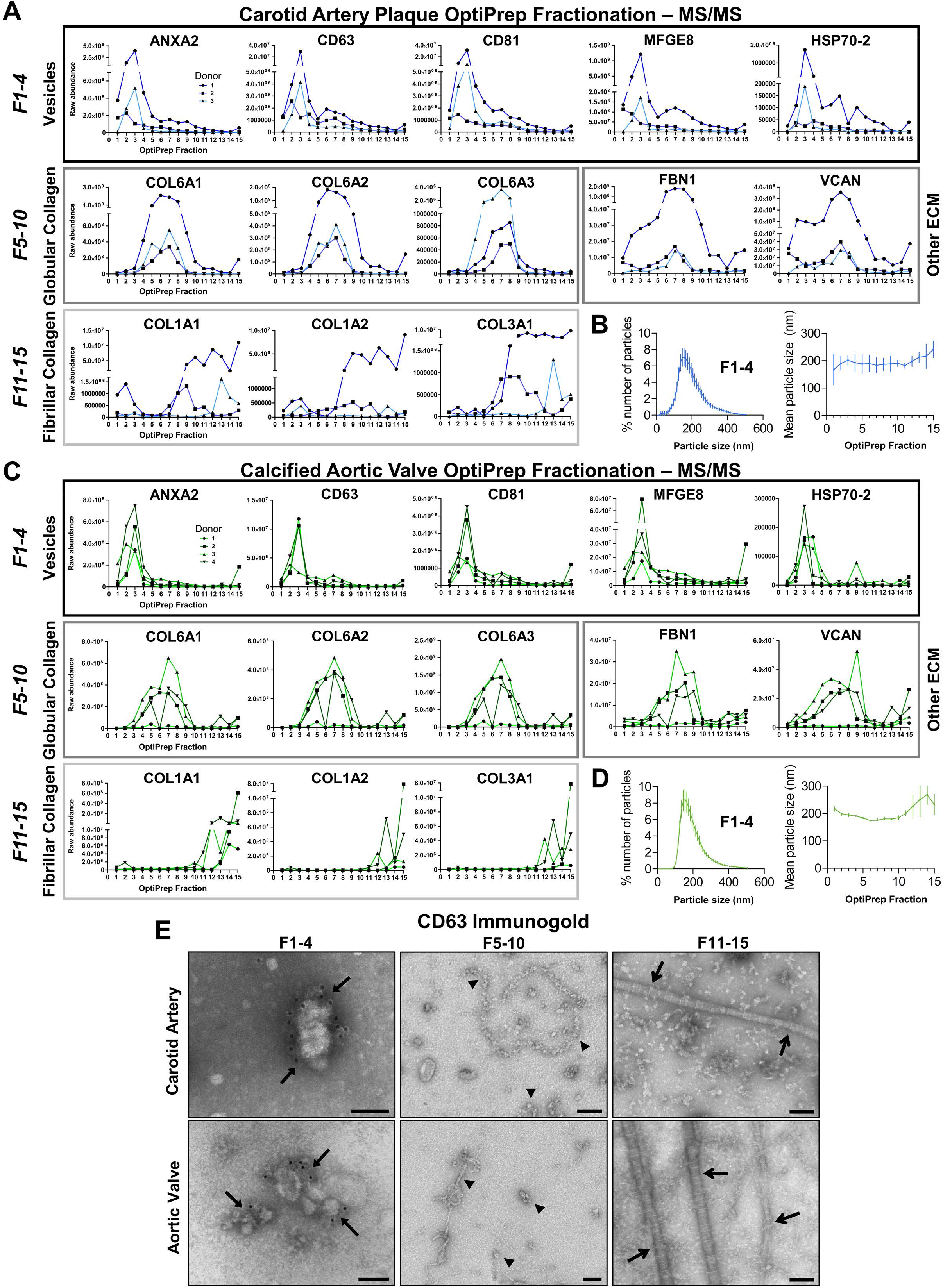
Tissue-Entrapped EVs are Highly Enriched in the Least-Dense Fractions from OptiPrep Density Gradient Separation. **A and C,** From 15-fraction survey experiments, raw abundance by mass spectrometry of selected proteins from human carotid artery plaques and calcified aortic valves demonstrated that extracellular vesicle marker proteins (Annexin A2, CD63, CD81, MFGE8 [lactadherin], HSP70-2) were highly enriched in the four least-dense fractions of both tissue types; n=3 carotid artery and 4 aortic valve donors. Enrichment of globular collagens (Collagen VIA1, VIA2, VIA3), fibrillar collagens (Collagen IA1, IA2, IIIA1), and other components of the extracellular matrix (ECM; FBN1, VCAN) was identified in denser OptiPrep fractions. **B and D,** Nanoparticle tracking analysis confirmed the characteristic presence of particles ~150-200 nm in diameter in OptiPrep fractions 1-4 of both carotid artery plaques and calcified aortic valves (n=3), while mean particle size remained consistent between OptiPrep fractions (mean±SEM). **E,** Representative CD63-labelled immunogold transmission electron microscopy (TEM) identified CD63+ membrane-bound EVs in OptiPrep fractions 1-4 (arrows) from carotid artery plaques (top) and calcified aortic valves (bottom); bar=100 nm. Consistent with mass spectrometry-derived protein abundance in A/C, TEM showed abundant globular collagens in fractions 5-10 (arrowheads) and fibrillar collagens in the most-dense fractions (open arrows).

Ultracentrifugation has frequently been used in isolation to enrich EVs from biofluids and *in vitro* cultures.(24) As tissues are inherently ECM-rich vs. 2D cell culture and biofluids, we performed mass spectrometry to compare calcified carotid artery and aortic valve tissue EVs enriched by our OptiPrep pipeline *vs.* ultracentrifugation alone (Supplemental Figure 5A). The OptiPrep-derived EV proteome was enlarged by nearly 40% vs. ultracentrifugation-alone in both vascular and valvular tissues, respectively (954 *vs.* 683 and 913 *vs.* 661 proteins; Supplemental Figure 5B). The proteome produced uniquely by ultracentrifugation was overwhelmingly composed of ECM and endoplasmic reticulum components (Supplemental Figure 5C, Supplemental Tables 7/8), while proteins found only after OptiPrep fractionation enrichment associated largely with intercellular-, membrane-, and vesicular components – fractions of the cell that are known to be sorted and loaded into EVs during vesicle biogenesis (Supplemental Table 9).(10) These OptiPrep-specific proteins also held diverse and physiologically-impactful functionalities (e.g. phosphorylation, protein transport, post-translational modification, and actin-binding; Supplemental Table 10) that may contribute to cell-cell communication. Lastly, we identified an effect of the iodixanol used in the OptiPrep density gradient on proteome depth and developed a mass spectrometric targeted mass exclusion strategy that improved the size of the OptiPrep-derived tissue EV proteome by 9.5% and contributed to further detection of pathologically-relevant EV functionality (Supplemental Figures 6, 7).

### Tissue-Entrapped EV Protein Cargoes in the Calcifying Vascular and Valvular Niches

EVs carry bioactive protein cargoes that can mediate intercellular signaling and act as early nucleation sites of microcalcification.(8) Having developed a pipeline for isolation and enrichment of tissue-entrapped EVs, we applied this toolkit to study whether tissue-specific alterations of entrapped EV cargoes exist in cardiovascular disease and examined putative contributions of EVs to the pathogenesis of vascular and valvular calcification.

To this end, we extracted tissue EVs from diseased vascular and valvular samples by pooling OptiPrep fractions 1-4 from an additional cohort of donors and confirmed EV enrichment and purification by TEM and NTA (Figure 4A). We observed a slight but significant increase in mean particle size of carotid artery *vs.* aortic valve EVs (230.7±3.3 *vs.* 193.5±5.1 nm, p<0.001, Figure 4A) that we have previously shown to correlate with degree of EV aggregation and calcification in 3D tissue culture.(8) Proteomics on these pooled fractions quantified 1,104 EV proteins, of which 96% were common to EVs from both tissue types (Figure 4B). To examine how EV cargoes reflect tissue-level processes amid disease, we assessed overlap of the whole-tissue and tissue-entrapped EV proteomes (Figure 4B): of 2,010 proteins detected, 769 (38.3%) were found in both the whole-tissue and tissue EV datasets. Correlation analyses of protein abundances for these shared proteins identified a moderately-strong Pearson’s r of 0.583 (p<0.0001), implying that the whole-tissue and EV proteomes are significantly correlated. EVs are thus reflective of some, but not all, phenotypes found at the whole-tissue level and appear to have important biological functions that could be missed by whole-tissue ‘omics (and vice-versa). Despite the large overlap in the complete EV proteome between tissue types, principal component and heat map analyses displayed strong tissue-specific clustering indicative of altered protein abundances therein (Figure 4B,C). In accordance with this finding, enrichment analysis identified 179 proteins that were statistically significantly differentially enriched between tissue types (114 up in calcified carotid artery EVs, 65 up in aortic valve EVs; FDR≤0.05; Figure 4C). To obtain biological insights from the proteomics data, we examined the contents of the tissue EV proteome through pathway enrichment analyses. In total, 442 significantly enriched KEGG, BioCarta, or Reactome pathways were obtained from the common EV proteome (1,063 proteins), 113 pathways from the 114 proteins that were enriched in carotid EVs, and 40 from the 65 proteins enriched in aortic valve EVs (Figure 4D, Supplemental Tables 11-13). Pathways linked to the common tissue EV proteome described a host of functions previously implicated in cardiovascular calcification, including vesicle transport/cell-cell communication, modulation of oxidative stress, ECM degradation, retinoid metabolism, and regulation of IGF transport/uptake. Interestingly, carotid artery EVs were specifically tied to pathways involved in responses to elevated calcium, platelet activation, inflammation, and mechanotransduction (including integrin binding, focal adhesions, cytoskeletal regulation, MAPK activation, and muscle contraction). In contrast, proteins enriched in aortic valve-derived EVs associated with a number of pathways involving cell cycle regulation by the tumor suppressor protein p53, as well as iron uptake, mineral absorption, sugar metabolism, and GLUT4 translocation.

**Figure 4:**
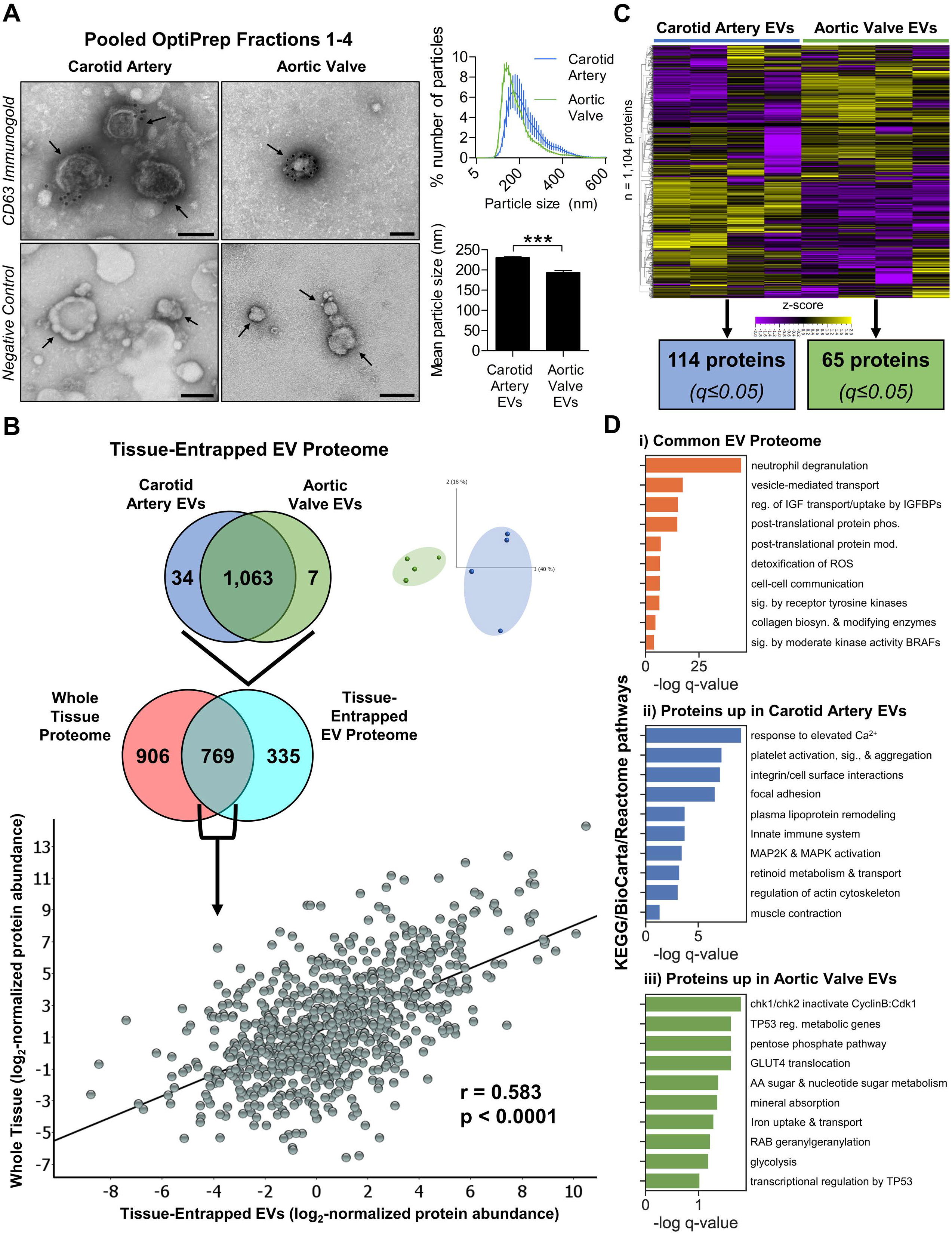
Proteomics of EV-Enriched Pooled Fractions to Quantify EV Protein Cargoes in Vascular and Valvular Tissue. **A,** Representative CD63-labelled immunogold transmission electron microscopy (TEM) and negative control images from pooled OptiPrep fractions 1-4 of human carotid artery plaque (left) and calcified aortic valve (right) demonstrated EV enrichment and purification; bar=100 nm. Nanoparticle tracking analysis found that EVs from carotid artery plaques had significantly larger mean diameters than those from calcified aortic valves (230.7±3.3 vs. 193.5±5.1 nm). **B,** The complete tissue-entrapped EV proteome was composed of 1,104 proteins. The carotid and valvular EV proteomes largely overlapped (1,063 EV proteins in common), though 41 proteins were uniquely detected in EVs from a single tissue type and principal component analysis identified tissue-specific clustering. 69.7% of the complete tissue-entrapped EV proteome (769 proteins) was also found in the whole-tissue proteome; linear regression identified a moderate and statistically significant correlation (Pearson’s r = 0.583, p<0.0001) of per-protein abundances between the tissue-entrapped EV and whole tissue proteomes. **C,** Heat map analyses (ordered by hierarchal clustering) of the complete tissue-entrapped EV proteome demonstrated alterations in individual EV protein abundances between cardiovascular tissue types. Enrichment analysis identified 179 EV proteins that were significantly differentially enriched between vascular and valvular EVs (proteins filtered at q≤0.05). **D,** Selected KEGG, Reactome, and BioCarta pathways that were significantly enriched in i) the common tissue-derived EV proteome, ii) proteins significantly enriched in carotid artery-derived EVs, and iii) proteins significantly enriched in aortic valve-derived EVs. Mean±SEM; *p<0.05, ***p<0.001; n=4 carotid artery and 4 aortic valve donors.

### Sequencing Reveals the Tissue-Derived EV Non-Coding miRNAome

Along with protein cargoes, noncoding miRs are also incorporated into EVs and can impact post-transcriptional regulation in recipient cells.(11) We extracted miRNA from the same OptiPrep-enriched diseased tissue EV samples that underwent proteomics (reported in Figure 4) and performed small RNA sequencing to assess the cardiovascular tissue-entrapped EV miRNAome (set of expressed microRNAs). RNA fragment analysis found that vascular- and valvular-derived EVs contained similar levels of microRNA (59.0±1.2 vs. 56.8±2.1% of all small RNA, p>0.05, Figure 5A), and 123 EV-derived miRs were mapped to miRbase after sequencing, of which 83% were common to calcified carotid artery and aortic valve EVs (Figure 5B, top). After annotation, miR counts were normalized, transformed, and tested for differential enrichment with DESeq2. The complete EV miRNAome clustered by tissue-type (Figure 5B, bottom) and 5 miRs were significantly differentially enriched between tissue types (2 up in carotid artery EVs, 3 up in aortic valve EVs; FDR≤0.05; Figure 5C): miR-143-3p, miR-145-5p, let-7c-5p, miR-125b-5p, and miR-574-3p. To determine the biological roles of the EV miRNAome, we used TargetScan 7.2 to predict high-confidence mRNA targets (≥95^th^ percentile weighted context++ score). The complete EV miRNAome associated with 5,182 high-confidence gene targets, of which miRs enriched in vascular and valvular EVs targeted 281 and 447 genes, respectively. We then built EV miRNA/mRNA target regulatory networks (Figure 5D) which exhibited clear clustering and segregation of target genes by tissue type, and as with the EV proteome, leveraged pathway enrichment analysis of EV miR gene targets to examine the functional impact of tissue-entrapped EV contents (Figure 5E). The target genes of the common tissue-entrapped EV miRNAome were significantly enriched among 677 pathways (Supplemental Table 14), including those involved in AGE-RAGE, mTOR, Hippo, FGFR, and Wnt signaling. miRs up in carotid artery EVs significantly targeted the constituents of 118 pathways (Supplemental Table 15), including several pathways associated with regulation of calcification and osteogenesis by the transcription factor RUNX2 and inflammatory toll-like receptor (TLR)/NF-κB signaling, while maintaining targeting of focal adhesion, cytoskeletal regulation, and muscle contraction pathways that were also tied to the carotid artery specimen EV proteome. Meanwhile, targets of miRs up in aortic valve EVs were significantly enriched in 155 pathways (Supplemental Table 16) and continued to target a multitude of p53-linked regulatory programs that were enriched in the aortic valve EV proteome, in addition to others involved in IL-17, sphingolipid, and PI3K-Akt signaling that play complementary roles in inflammation and calcification.

**Figure 5:**
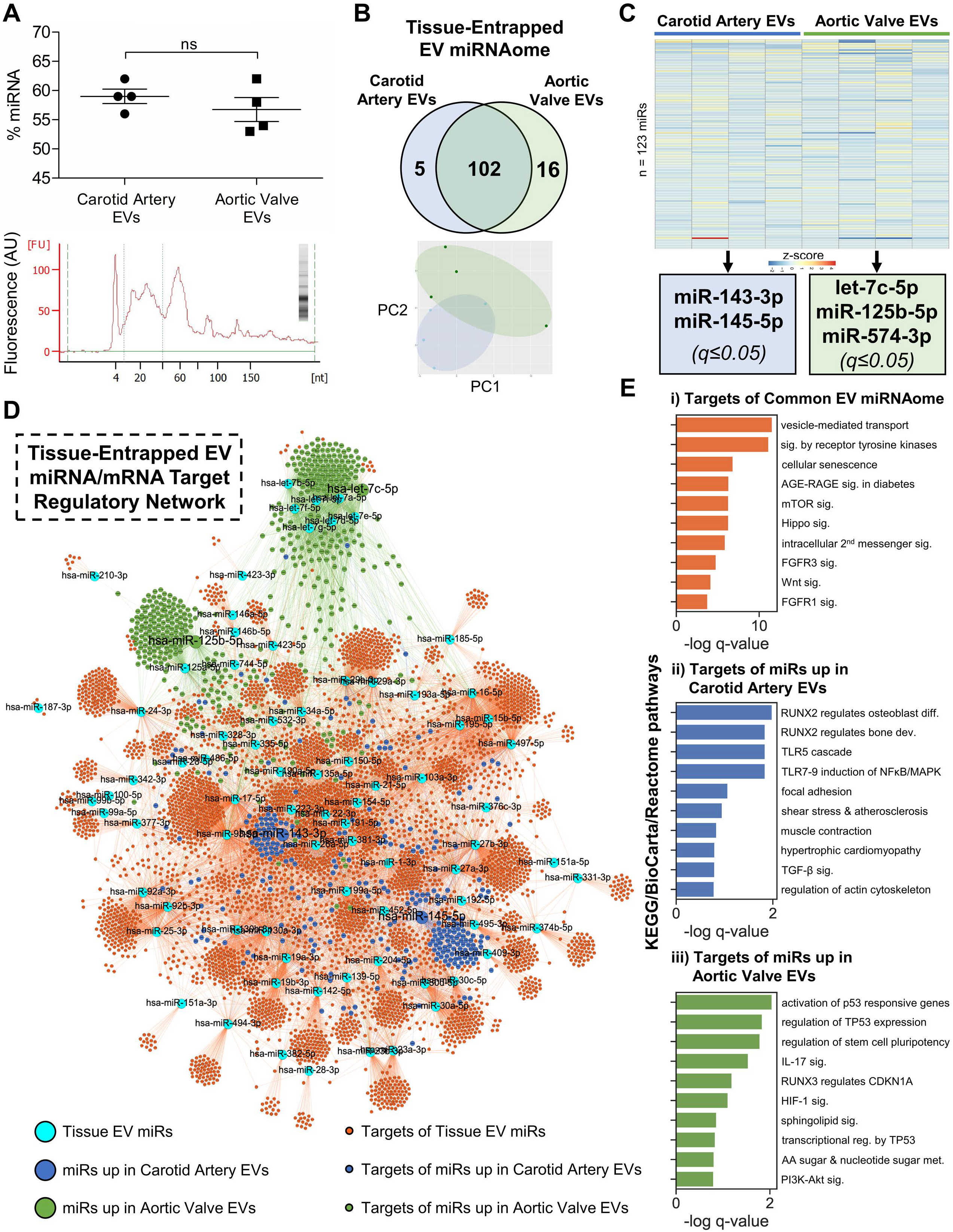
Sequencing Reveals the Human Tissue-Entrapped EV non-coding miRNAome from Vessels and Valves. **A,** RNA fragment analysis found that OptiPrep-enriched EVs contained enhanced levels of miRNA (mean miRNA content=57.3% of all small RNA). miRNA enrichment was unchanged between calcified carotid artery plaque- and aortic valve-derived EVs (top). Bottom: representative fragment analysis tracing showing large miRNA-associated fragment peak at 24 nucleotides (nt). **B,** Of 123 miRs sequenced in the complete tissue-entrapped EV miRNAome, 21 were found exclusively in EVs extracted from one cardiovascular tissue type, while 102 were common to both vascular and valvular EVs; principal component analysis identified tissue-specific clustering of EV miRs. **C,** Though levels of individual EV miRs were largely unchanged between cardiovascular tissue types as demonstrated by heat map analysis (ordered by q value), 5 miRs were significantly differentially enriched between vascular and valvular EVs (miR-143-3p, miR-145-5p, let-7c-5p, miR-125b-5p, miR-574-3p; q≤0.05). **D,** miRNA/mRNA target gene regulatory networks of 5,182 TargetScan-predicted gene targets (≥95^th^ percentile weighted context++ score) of the complete tissue-derived EV miRNAome exhibited clear clustering and segregation of miR target genes by tissue type. **E,** Selected KEGG, Reactome, and BioCarta pathways significantly enriched in i) the 4,765 target genes of the common tissue-derived EV miRNAome, ii) 281 target genes of miRs enriched in carotid artery-derived EVs, and iii) 447 target genes of miRs increased in aortic valve-derived EVs. Mean±SEM; n=4 carotid artery and 4 aortic valve donors.

### Multi-omics Integration Identifies Conserved Functionality of Cardiovascular Tissue-Entrapped EV Cargoes

While the protein and noncoding miR contents of EVs are both bioactive and known to modulate recipient cell phenotype and function, their impact has, to our knowledge, previously only been studied separately from one another. To assess the unified effect of these two types of cargoes, we completed pathway-level integration on multiple layers of EV omics data. We began by integrating cargoes common to EVs from both calcified carotid artery plaques and aortic valves, as these cargoes represent putatively conserved cardiovascular tissue EV functionality. 154 pathways (a significant overlap, p<0.001; Supplemental Table 17) were enriched in both the common EV proteome and gene targets of the common EV miRNAome (Figure 6A, left). To dissect the conserved biological impact of these 154 shared pathways, we assembled them into a pathways network, based on the Jaccard index of overlap between pathway members that were detected in our proteomics and transcriptomics (Figure 6A, right). Unbiased Louvain clustering (25) was then used to identify functional communities within the pathway network, revealing that EV cargoes from diseased cardiovascular tissues shared 9 distinct functions including regulation of the cell cycle, synthesis and organization of ECM, and modulation of MAPK and Rho GTPase intracellular signaling cascades (Figure 6B, Supplemental Figure 8, Supplemental Table 18). Together, these data indicate that cardiovascular tissue-derived EVs carry a potent ability to modulate intercellular signaling and recipient cell phenotype in disease.

**Figure 6:**
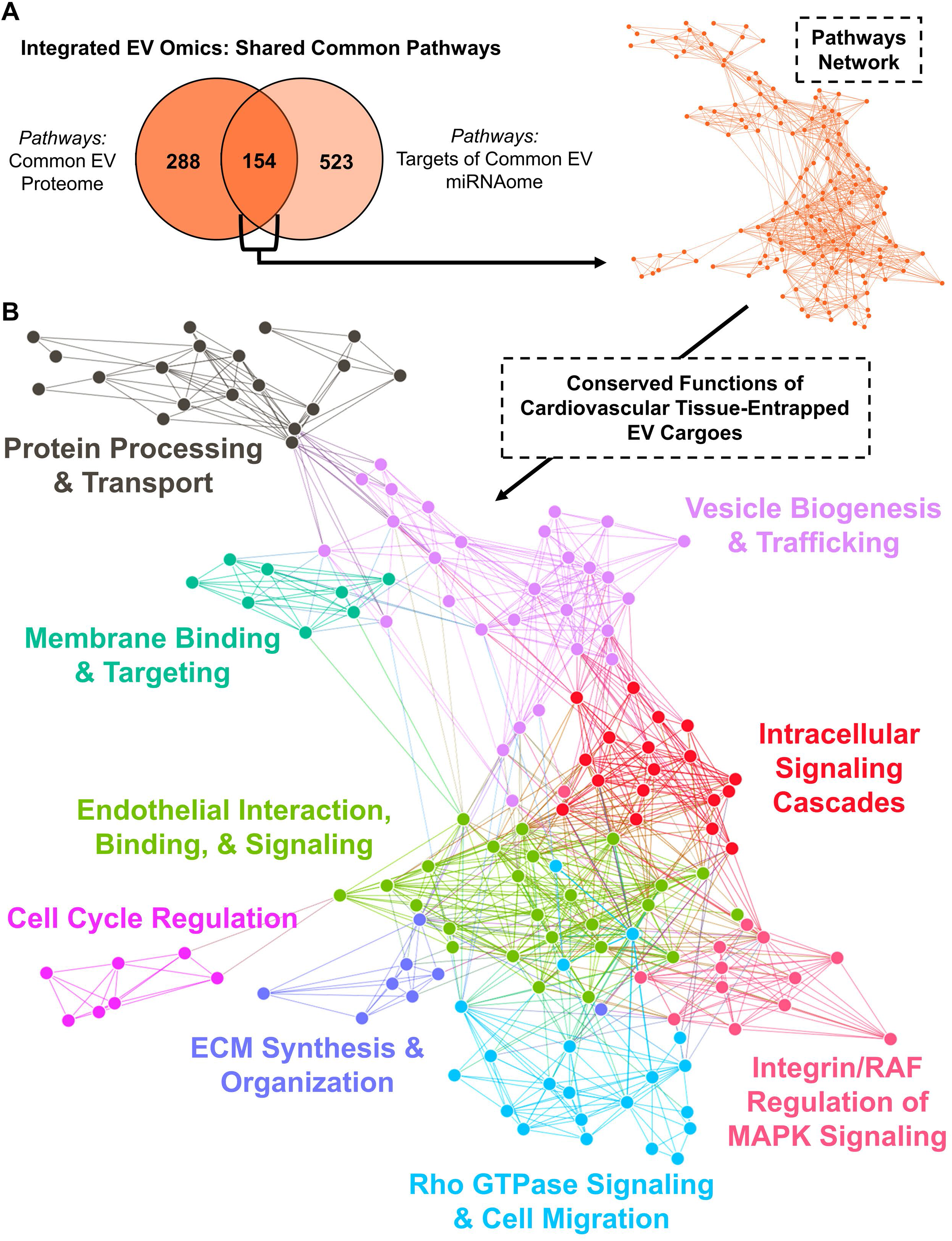
Pathway Networks Shared Across Omics Layers Identify Conserved Functionality of Cardiovascular Tissue-Entrapped EV Cargoes. **A,** A statistically significant number of KEGG, Reactome, and BioCarta pathways (154 pathways, p=0.000206) were significantly enriched in both the proteome and gene targets of the miRNAome that was common to calcified carotid artery and aortic valve-derived EVs (n=4). The network of these overlapping pathways was generated with pathways as the nodes (node size corresponds to −log(q-value)) and shared detected genes between pathways as the edges (edge thickness matches the Jaccard index of overlap between detected genes of the two connected pathway nodes). **B,** Unbiased clustering of pathways into real network communities by the Louvain method revealed 9 distinct functions shared by cardiovascular tissue-derived EV cargoes, including cell cycle regulation, synthesis and organization of the extracellular matrix (ECM), and modulation of MAPK and Rho GTPase intracellular signaling cascades.

### Network Analyses Across Omics Layers Implicate Mechanotransduction and p53 Transcriptional Regulation in the Tissue-Specific Effects of Cardiovascular EVs

Though we found substantial overlap of the detected proteome (96%) and miRNAome (83%) between vascular and valvular tissue EVs, clustering and differential enrichment analyses (Figures 4B,C and 5B,C) indicated that a subset of protein and miR cargoes were altered amongst these tissue types. We therefore also performed tissue-specific multiomics integration of EV cargoes by evaluating the overlap of pathways that were significantly associated with (i) proteins and (ii) gene targets of miRs that were differentially enriched in (iii) carotid artery- or (iv) aortic valve-derived EVs (Figure 7A, Supplemental Table 19). Eight pathways were shared by both the protein and miR targets within a tissue type (6 in calcified carotid artery and 2 in aortic valve-derived EVs), indicating shared tissue-specific bioactivity of EV contents. To visualize the integrated impact of these shared EV contents, we constructed pathway-specific protein-protein interaction networks by mapping pathway genes onto the STRING PPI network, then highlighting those constituents of the network that were significantly enriched in the EV proteome or miRNAome (Figure 7B,C, Supplemental Figures 9-14). Along with pathways associated with focal adhesions, actin cytoskeleton regulation, and cardiomyopathies, constituents of the Reactome “Muscle Contraction” pathway (R-HSA-397014, Figure 7B) were significantly enriched in multiple layers of calcified carotid artery EV omics. This finding hints further at a shared role for vascular EV cargoes in regulating cellular adhesion and migration *in vivo* that has previously been demonstrated in the proteome of EVs from cultured vascular SMCs,(15) and ample evidence indicates that elevated intracellular calcium levels modulate proteins involved in vascular SMC mechanotransduction and resultant fibrocalcific differentiation. In addition to a pathway associated with amino and nucleotide sugar metabolism, the “Transcriptional Regulation by TP53” pathway (R-HAS-3700989, Figure 7C) was significantly enriched in components from multiple layers of aortic valve EV omics; modulation of p53 has recently been strongly implicated in cardiovascular calcification by impacting NOTCH1, *Slug,* and BMP2 signaling.(22, 26–28)

**Figure 7:**
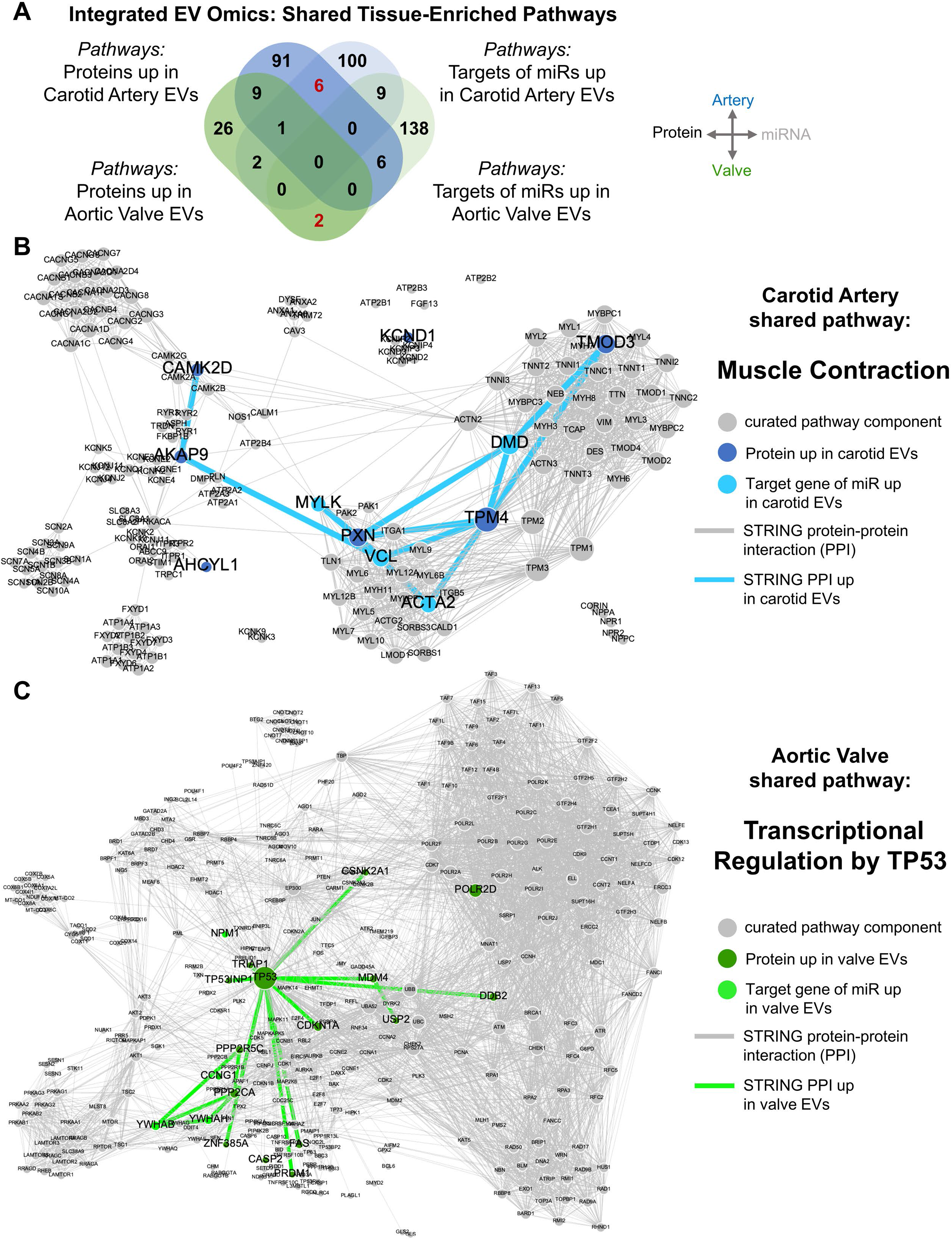
Integration of EV Multi-Omics Identifies Tissue-Specific EV Functions. **A,** Overlap of KEGG, Reactome, and BioCarta pathways significantly associated with proteins and gene targets of miRs that were differentially enriched between calcified carotid artery and aortic valve-derived EVs (n=4). No pathways were shared between all four groups (proteins *and* miR targets in artery *and* valve-derived EVs). However, several pathways were shared by the proteins and targets of miRs within a particular tissue (6 and 2 in artery- and valve-derived EVs, respectively; in red). **B and C,** Pathway-specific protein-protein interaction networks were utilized to integrate EV multi-omics. Along with pathways associated with focal adhesions, actin cytoskeleton regulation, and cardiomyopathy, constituents (blue nodes) of the Reactome “Muscle Contraction” pathway (R-HSA-397014) were significantly enriched in multiple layers of carotid artery-derived EV omics. In addition to a pathway associated with amino and nucleotide sugar metabolism, the “Transcriptional Regulation by TP53” pathway (R-HAS-3700989) was significantly enriched in components (green nodes) from multiple layers of aortic valve-derived EV omics. Node diameter corresponds to node degree.

## Discussion

EVs regulate a wide spectrum of key biological processes, act as mediators of cell-cell communication,(11) predict cardiovascular events,(12) and actively form the building blocks of cardiovascular microcalcification.(8) This work represents, to the best of our knowledge, the first time that EV cargoes have been studied from tissues other than easy-to-dissociate brain (17) and cell-line-derived mouse adenocarcinoma.(18) Unlike in biofluids or brain, extraction of EVs from atherosclerotic plaques or diseased aortic valves is particularly challenging due to the extensive ECM-rich fibrosis and severe calcific burden present in these tissues upon excision. Loss of cellular integrity due to mechanical tissue homogenization leads to co-isolation of intracellular vesicles or artifactual membranous particles,(29) and the apoptotic and necrotic nature of these diseases can also leave the extracellular microenvironment replete with large apoptotic bodies. Previous attempts to extract large microparticles from atherosclerotic human plaques by mincing and centrifugation alone (30, 31) or from calcified aorta by tissue grinding, enzymatic digestion, and ultracentrifugation (15) were prone to these potential sources of contamination. Gentle enzymatic digestion using the protease papain has been implemented on brain,(18) though these studies also required Dounce homogenization because of papain’s limited ability to digest collagen. Our use of bacterial collagenase negated the need for mechanical homogenization (avoiding collection of intracellular vesicles or artifactual particles) and enabled digestion of collagen-rich cardiovascular tissues, maintenance of cell integrity/viability, and effective *in silico* elimination of the enzymatic protein. This approach allowed co-isolation of both viable primary cells for tissue culture and EVs from the same piece of tissue, thus permitting *in vitro* counterpart experiments. By combining ultracentrifugation with OptiPrep density gradient separation, we demonstrate enrichment of EVs from cardiovascular tissues and thoroughly classify them based on their size, density, morphology, and protein/nucleic acid cargoes in accordance with the MISEV2018 guidelines.(29) Our approach reveals a dramatically-improved ability to remove co-isolated non-EV contaminants that are present should tissue digests solely undergo high-speed and/or ultracentrifugation prior to downstream analyses.(15) We also show that enrichment of EVs from ECM-rich tissues by size exclusion chromatography may be difficult due to the robust presence of collagen fragments in the same size range as EVs – differing densities and molecular weights instead allow successful separation by OptiPrep gradients. The use of iodixanol alone in our density gradient is key, as prior studies in brain have relied on sucrose-based gradients, which can cause leaching of vesicle cargoes during isolation.(32) Here we describe a means to account for the impact of iodixanol on mass spectrometric peptide identification.

A common set of risk factors predispose patients to cardiovascular calcification, however only a subset of patients suffer from calcification of both the vessels and valves.(2) Our direct comparison of whole-tissue proteomes revealed novel insights into tissue-specific modes of pathogenesis and identified a set of highly differentially-expressed proteins that have potential usage as biomarkers to discriminate between these two pathologies. In this manner, we revealed specific enrichment of proteins and pathways involved in cell adhesion, migration, and cytoskeletal force generation, inflammation, and lipid metabolism in calcified carotid artery plaque. Our whole-tissue results also imply that calcific aortic valve disease is over-represented in aspects of osteochondral and proteoglycan-associated components when compared with that of atherosclerotic plaques; these features are hallmarks of early valvular stenosis in swine and humans.(3) Electron microscopy has shown calcified EVs to be present in mineralized regions of both tissues,(8, 16) consistent with the substantial enrichment of vesicle ontologies that we found to be shared by the proteomes of carotid artery plaque and calcified aortic valves at the whole-tissue level. Overall, the cardiovascular EV proteome and miRNAome were associated with pathways that included intercellular vesicle-mediated communication and a range of functions that are broadly incriminated in cardiovascular calcification: collagen metabolism, receptor tyrosine kinase signaling, Wnt signaling, FGFR signaling, oxidative stress, and insulin-like growth factor transport. Our data suggest that EV cargoes act as cell-cell mediators of these pathological processes within both the atherosclerotic plaque and calcific valve. In the context of vascular *vs.* valvular calcification, roughly 16% of the EV proteome was differentially enriched between tissue types, suggesting that this handful of proteins delineate pathophysiology of these two disease entities. The carotid artery EV proteome linked to mechanotransduction, calcium response, and retinoid metabolism pathways. In contrast, EVs from calcified aortic valves associated directly with pathways including mineral absorption and iron binding capability. Of note, iron uptake by VICs after interleaflet hemorrhage has recently been shown to induce VIC proliferation, osteogenic differentiation, and calcification.(33)

We also identified tissue-specific enrichment of a select set of EV miRs, some of which have been previously implicated in cardiovascular calcification, though not in the context of EV-mediated delivery. Levels of the miR-143/145 cluster are elevated in OptiPrep-enriched carotid artery-derived EVs; miR-145 promotes differentiation of SMCs to contractile phenotypes by targeting KLF4 signaling(34), is increased in symptomatic *vs.* asymptomatic atherosclerotic plaques, and correlates with plaque destabilization.(35) miRs enriched in valvular EVs included miR-125b-5p; whose modulation drives osteoblastic differentiation in vascular SMCs and whose levels are altered in aged *Apoe*^−/−^ mice fed a high-fat diet.(36) miR-125b is also a critical regulator and driver of cardiac fibrosis, as it induces myofibroblastic transition via apelin inhibition and drives fibroblast proliferation via p53.(37) Therefore, valvular EV miR-125b-5p may modulate the expansion and differentiation of VICs down a pathogenic fibrocalcific lineage. The other miRs upregulated in valvular EVs are reported to exert control over osteochondrogenesis: let-7c-5p promotes osteogenic differentiation and bone formation by mesenchymal stem cells (MSCs) in response to inflammation through interactions with the HMGA2/PI3K/Akt signaling cascade,(38) while miR-574-3p is a mechanoresponsive miR (particularly relevant to the biomechanically-complex valvular microenvironment) known to regulate MSC chondrogenesis downstream of Sox9 via targeting of the retinoid X receptor (RXR)α.(39) Pathway analysis on the predicted targets of these miRs strongly implicated RUNX2, toll-like receptors, and mechanotransduction as key downstream pathological effectors of vascular EVs, while valvular EV miRs were linked to IL-17 and sphingolipid signaling pathways that are involved in valvular inflammatory cell infiltration and osteogenesis/calcification.(40)

Our findings suggest that advanced cardiovascular disease takes a largely similar form in both atherosclerotic plaque and calcified aortic valve: 99.3% of the whole-tissue proteome was identified in both tissue types, along with 96.3% of the tissue EV proteome, 82.9% of the tissue EV miRNAome, and 92.0% of gene targets of the EV miRNAome. Nevertheless, there are remarkably consistent tissue-specific alterations in protein/miR abundance that indicate divergent pathogenesis, which could be exploited for tissue-specific therapeutics: at the whole-tissue level, 15.9% of proteins are differentially enriched between carotid artery plaque and calcified aortic valve, along with 16.2% of the tissue EV proteome. While only 4.1% of the tissue EV miRNAome is differentially enriched, gene targets of differentially enriched miRs represented 14.0% of all tissue EV miRNAome gene targets. A strength of our approach is that integration of these multiple omics layers allows for coherent biological insight to be drawn from the perspective provided by any one omics modality.(21) To that end, multi-omics integration offered the first means to examine the combined impact of protein and RNA EV cargoes on recipient cells, the plurality of which was composed of pathways associated with intracellular signaling cascades. Overall, EV cargoes were connected to RAF regulation of MAPK signaling (control of proliferation and mechanosensitive osteogenesis)(41) and Rho GTPase-associated regulation of cellular migration.(42) In addition, integration of carotid artery-derived EV protein and miR contents revealed a strong tissue-specific signal for mechanotransduction-mediating pathways involving focal adhesions, integrin binding, regulation of actin organization, and contractile force production that are well-understood to regulate cell adhesion and migration. Phenotypic switching of vascular SMCs from a contractile to a proliferative/migratory state contributes to both initial plaque development and fibrous cap formation (reviewed in (43)), and our data suggest that EVs help to drive this transition. Notably, proteins found in EVs from cultured vascular SMCs have previously been hypothesized to regulate cellular adhesion and migration(15), perhaps in response to vascular injury. We show that this effect is also present *in vivo*, is preferentially enriched in calcified atherosclerotic plaques when compared to calcified valves, and we expand this regulatory paradigm to include the impact of the carotid EV miRNAome. We also identified a robust link between aortic valve-derived EV cargoes and functional pathways involving the transcription factor p53. Reductions in p53 recruitment due to upregulation of the lncRNA H19 reduce Notch signaling and induce VIC mineralization through RUNX2 and BMP2 signaling.(26) In hyperphosphatemic calcification, p53 levels correlate inversely with osteogenic differentiation and calcific burden in SMC culture and in mice with chronic kidney disease.(28) Others have shown that inhibition of vascular calcification by tenopiside occurs via the BMP2/RUNX2 axis in a p53-dependent manner – inhibition of BMP2 expression (and thus of calcification) was linked with activated p53. Directionality of p53 regulation in valvular calcification remains complicated, as warfarin treatment induces p53 overexpression and calcification in VIC cultures and murine aortic valves via recruitment to the *Slug* promoter.(27) While further studies will be needed to delineate the mechanistic contributions of valvular EVs to p53 transcriptional regulation, our data determined a putative role for EVs in contributing to this aspect of calcific valve disease.

## Conclusions

The present study compares and contrasts the proteomes of human calcific vascular and valvular disease. After having identified elevated cellular contractility and inflammation in carotid artery plaque along with increased osteochondral disease signatures in the aortic valve, we found a significant and shared presence of vesicle-associated functionality at the whole-tissue level. We then delineate a combination of enzymatic digestion, (ultra)centrifugation, and density gradient separation to rigorously enrich EVs from calcified human atherosclerotic plaques and aortic valves, and interrogate both the protein and non-coding RNA contents of EVs that become entrapped in these cardiovascular tissues during disease progression. Those EV cargoes carry both conserved and tissue-specific pathogenic functionalities in vascular *vs.* valvular calcification and shed light on how common risk factors may drive divergent disease processes *in vivo*. We tied tissue-entrapped EV cargoes to tissue-specific regulation of two key pathogenic mechanisms: cellular adhesion/migration in carotid artery plaque, and p53-mediated transcriptional regulation in calcific aortic valve disease. This work opens an avenue towards further examination of the critical roles that vesicles play as mediators of cell-cell communication, cellular differentiation, and cell-matrix interactions in the circulatory system. Evidence suggests that both circulating and tissue-entrapped EV pools likely contain sub-populations of EVs that are secreted by different cell types, have exosomal or microvesicular origins, and function in protective or pathological ways.(44) Here, we take a step towards the examination of these EV populations and their complex interplay in cardiovascular calcification. Ultimately, our approach represents an efficient, cost-effective, and widely accessible strategy for studying the proteome and miRNAome of purified EVs isolated directly from fibro-calcific human, animal, or even engineered tissues.(8, 45)

## Materials and Methods

Unless otherwise noted, all materials were purchased from Sigma-Aldrich.

### Human tissue collection

Human aortic valve (AV) leaflets were obtained from AV replacement surgeries for AV stenosis (Brigham and Women’s Hospital IRB protocol #: 2011P001703), and atherosclerotic human carotid artery plaque samples were obtained from patients undergoing carotid endarterectomies (Brigham and Women’s Hospital IRB protocol #: 1999P001348) from a total of 27 donors. Fresh tissue was immediately transferred from the operating room in high-glucose Dulbecco’s modified Eagle’s medium (DMEM, Gibco 10569010) on ice and was stored at 4°C until further processing within 45 minutes of extraction in the operating room.

### Whole-tissue lysis

Calcified stenotic aortic valve and carotid plaque tissue samples were each dissected into non-diseased, fibrotic, and calcified macroscopically distinct segments per sample as described previously.(21) Tissue samples were then pulverized in liquid nitrogen and re-suspended on ice in RIPA buffer (Thermo Scientific Pierce, 89900) with protease inhibitor (Roche, cOmplete ULTRA tablets, 45892970001) and phosphatase inhibitor (Roche, PhosSTOP tablets, 4906845001), and vortexed for 30 seconds.

### Extracellular vesicle isolation

#### Enzymatic digestion

Valvular and carotid tissue samples underwent enzymatic digestion largely as described previously.(46) In brief, tissue samples were rinsed briefly in sterile phosphate buffered saline without calcium or magnesium (PBS−/−, Corning 46-013-CM), scraped gently on each side with a razor blade to remove the endothelium, and rinsed again. Tissue was then roughly chopped into ~5 mm^3^ cubes and incubated at 37°C for 1 hour in 1 mg/ml of filter-sterilized collagenase type IA-S from *Clostridium histolyticum* (Sigma-Aldrich C5894) in DMEM, mixing every 20 minutes. After 1 hour, samples were vortexed briefly, the collagenase solution was removed, and tissue pieces were rinsed briefly with DMEM. The collagenase solution and DMEM rinse were pooled, centrifuged at 500xg for 5 minutes, and the resulting supernatant was stored at 4°C. Tissue pieces were then incubated again in 1 mg/ml collagenase for a further 3 hours at 37°C, mixing every 30 minutes. After 3 hours, the digest was vortexed briefly and strained through a 40 μm cell strainer. A DMEM rinse followed and was again strained at 40 μm. The resultant collagenase solution and DMEM rinse were pooled and centrifuged at 500xg for 5 minutes. The resultant 3-hour supernatant was pooled with the 1-hour supernatant and stored at −80°C. The 500xg pellet was resuspended in growth medium (DMEM with 10% fetal bovine serum (FBS, Gibco), 100 units/ml penicillin, and 100 μg/ml streptomycin), seeded on uncoated tissue-culture-treated polystyrene (TCPS), and cultured at 37°C and 5% CO_2_. Media was changed every 3 days.

#### Ultracentrifugation

In 15-fraction survey or EV-enriched pooled fraction experiments, the 500xg supernatant from enzymatic digestion of cardiovascular tissues underwent further ultracentrifugation at 10,000xg for 10 minutes at 4°C (Beckman Coulter, Optima MAX-UP, fixed-angle rotor MLA-55) in 10 ml 16×76 mm polycarbonate uncapped centrifuge tubes (Beckman Coulter 355630). The resultant supernatant was collected and underwent ultracentrifugation at 100,000xg for 40 minutes at 4°C (rotor MLA-55) in 10.4 ml 16×76 mm polycarbonate capped centrifuge tubes (Beckman Coulter 355603), the pellet was washed once in PBS−/− and centrifuged again at 100,000xg for 40 minutes at 4°C. The final pellet was resuspended in 150 μl of NTE buffer (137 mM NaCl, 1 mM EDTA, 10 mM Tris, pH 7.4) with protease inhibitor (Roche, cOmplete Mini tablets, 4693159001). In ultracentrifugation library experiments, the 500xg supernatant was first spun at 10,000xg for 10 minutes at 4°C (Beckman Coulter, fixed angle rotor TLA 120.2) in 1 ml 11×34 mm polycarbonate uncapped centrifuge tubes (Beckman Coulter 343778), and the resultant supernatant was ultracentrifuged at 100,000xg for 40 minutes at 4°C (rotor TLA 120.2; Beckman Coulter 343778 1 ml uncapped tubes).

#### Density gradient centrifugation

In 15-fraction survey or EV-enriched pooled fraction experiments, re-suspended ultracentrifugation-derived pellets were then layered onto the top of a linear 5-step 10-30% iodixanol gradient (composed of NTE buffer and OptiPrep Density Gradient Medium, Sigma-Aldrich D1556) in 10.4 ml 16×76 mm polycarbonate capped centrifuge tubes (Beckman Coulter 355603). The iodixanol gradient was then ultracentrifuged at 250,000xg for 40 minutes at 4°C (MLA-55), and 15 fractions were collected from the top of the gradient. In 15-fraction survey experiments, each resultant fraction underwent separate ultracentrifugation at 100,000xg for 40 minutes at 4°C (rotor TLA 120.2) in 1 ml 11×34 mm polycarbonate uncapped centrifuge tubes (Beckman Coulter 343778). In EV-enriched pooled fraction experiments, fractions 1-4 were pooled together, topped up to a volume of 9 ml with NTE buffer, and underwent ultracentrifugation at 100,000xg for 40 minutes at 4°C (rotor MLA-55; Beckman Coulter 355603 10.4 ml capped tubes). Supernatant was discarded from each fraction, and the resultant pellets were re-suspended in buffers appropriate to their downstream applications (detailed below).

### Transmission electron microscopy

5 μl of EVs (from iodixanol fraction(s) of interest, immediately after the 250,000xg density gradient centrifugation) were adsorbed for 1 minute onto carbon-coated, standard thickness, 400 mesh support film grids (EMS CF400-CU) made hydrophilic by plasma treatment (glow discharge, 25 mA). Excess liquid was removed with Whatman grade 1 filter paper (Sigma-Aldrich WHA1001325), and grids were rinsed of phosphate and salts by floating briefly on a drop of water. For immunogold labelling, samples were blocked with 1% bovine serum albumin (BSA) for 10 minutes, and incubated with mouse anti-human CD63 primary antibody (BD Pharmingen 556019, 1:20 in 1% BSA) for 30 minutes at room temperature. Grids were washed with 3 drops of PBS−/− in 10 minutes, then stained with rabbit anti-mouse bridging antibody (Abcam ab6709, 1:50 in 1% BSA) for 10 minutes. After further rinsing, grids were incubated with 10 nm Protein A-gold particles (University Medical Center Utrecht,1:50 in 1% BSA) for 20 minutes. Samples were then washed in PBS−/− (2 changes in 5 minutes) and water (4 changes in 10 minutes). Grids were blotted again, then stained with 0.75% uranyl formate (EMS 22451) for 15 seconds. Excess uranyl formate was then removed by blotting, and grids were examined under a transmission electron microscope (JEOL 1200EX) and imaged with a CCD camera (AMT 2K).

### Nanoparticle tracking analysis

Particle size and concentration was measured by nanoparticle tracking analysis (NTA, Malvern Instruments, NanoSight LM10). Before injection into the laser-illuminated chamber, samples were diluted 1:100 in PBS−/− (to ~10^9^ particles/ml). For each sample, five data collection windows (1 minute per window) were recorded during continuous injection by a syringe pump (Malvern Instruments). Particles were detected and quantified at screen gain=1.0, camera level=9.0 for capture and screen gain=10.0 and detection threshold=2.0 for processing. Particle counts presented are the average of 5 collection windows. Particle size distributions are reported as the mean of all donors after sum normalization per donor. Mean particle size is reported as the mean of all donors after averaging all particles per donor or per fraction per donor. Data are presented as mean ± standard error, and Student’s *t*-test (two-tailed, unpaired) was used for statistical comparisons between tissue types.

### Protein quantification

In experiments where input to trypsin/RapiGest proteolysis or to the iST kit were normalized by protein amount (see below), protein yield was quantified by the bicinchoninic acid assay (BCA, Pierce 23225) on a NanoDrop 2000 spectrophotometer (ThermoFisher Scientific) reading at 562 nm against a pre-mixed standard curve.

### Proteomics sample preparation

To obtain peptides for mass spectrometry, whole-tissue samples were sonicated after RIPA lysis for 4 × 15 seconds (Branson Sonifier 450). Protein precipitation was performed by methanol-chloroform and proteolysis by trypsin/RapiGest (Promega Gold Grade, V5280/Waters RapiGest SF, 186001861) as described previously.(47) 15 μg of precipitated protein per sample was used for proteolysis. Tryptic peptides were desalted with Oasis HLB 1 cc/10 mg cartridges (Waters 186000383) and dried with a tabletop speed vacuum (Thermo Scientific SPD1010), then re-suspended in 40 μl of 5% mass spectrometry grade acetonitrile (Thermo Fisher Scientific) and 0.5% formic acid. EV samples were processed with the PreOmics iST kit (PreOmics GmbH, P.O.00027) according to the manufacturer’s recommended protocol (v2.6), without sonication and with a 1.5-hour incubation at 37°C. Input to the iST kit varied by sample type. For 15-fraction survey experiments, the pellet from each fraction was resuspended in 12 μl of LYSE, and 10 μl were loaded into the iST kit. EV pellets obtained by ultracentrifugation alone were each re-suspended in 12 ul of LYSE lysis buffer (PreOmics); 2 μl of the resultant solution was utilized for protein quantification by BCA, and a volume of LYSE equivalent to 5 μg of protein was loaded into the iST kit and topped up to 10 μl with additional LYSE. For EV-enriched pooled fractions, the pellet resulting from ultracentrifugation of pooled fractions 1-4 was re-suspended in 12 μl of LYSE; 2 μl of the resultant solution was utilized for protein quantification by BCA, and a volume of LYSE equivalent to 5 μg of protein was loaded into the iST kit and topped up to 10 μl with additional LYSE. In all cases, peptides produced by the iST kit were resuspended in 40 μl of LC-LOAD (PreOmics).

### Mass spectrometry

Peptide samples were analyzed on an Orbitrap Fusion Lumos mass spectrometer fronted with an EASY-Spray Source (heated at 45°C), and coupled to an Easy-nLC1000 HPLC pump (Thermo Scientific). The peptides were subjected to a dual column set-up: an Acclaim PepMap RSLC C18 trap analytical column, 75 μm X 20 mm (pre-column), and an EASY-Spray LC column, 75 μm X 250 mm (Thermo Fisher Scientific). The analytical gradient was run at 300 nl/min, with Solvent A composed of water/0.1% formic acid and Solvent B composed of acetonitrile/0.1% formic acid). The acetonitrile and water were LC-MS-grade. For 30-minute gradients (ultracentrifugation libraries, stand-alone 15-fraction surveys), the analytical gradient was run from 5-21% Solvent B for 25 minutes and 21-30% Solvent B for 5 minutes. For 90-minute gradients (whole-tissue samples, ultracentrifugation libraries, EV-enriched pooled fractions), the analytical gradient was run from 5-21% Solvent B for 75 minutes and 21-30% Solvent B for 15 minutes. Whole-tissue samples (15 μg input) were diluted 1:20 in loading buffer prior to injection. Stand-alone 15-fraction surveys were injected at 1X. Ultracentrifugation libraries (5 μg input) and EV-enriched pooled fractions (5 μg input) were diluted 1:2 in loading buffer prior to injection. The Orbitrap analyzer was set to 120 K resolution, and the top N precursor ions in 3 seconds cycle time within a scan range of 375-1500 m/z (60 seconds dynamic exclusion enabled) were subjected to collision induced dissociation (CID; collision energy, 30%; isolation window, 1.6 m/z; AGC target, 1.0 e4). The ion trap analyzer was set to a rapid scan rate for peptide sequencing (MS/MS). When targeted mass exclusion of iodixanol was performed, it was enacted at an m/z of 775.8645 (z=2) with an exclusion mass width of 10 ppm. The retention time window was determined by pilot injections for each sample, with a 4-minute retention time window of beginning between 17-18 minutes for the 30-minute gradients and a 7-minute window starting at 28 minutes for the 90-minute gradient.

Resultant whole-tissue MS/MS data were queried against the Human (UP000005640, downloaded November 21, 2018) UniProt database, and tissue-derived EV MS/MS data were queried against the Human (UP000005640, downloaded July 7, 2014 for stand-alone 15-fraction survey experiments or November 21, 2018 for all others), *Hathewaya histolytica* (May 16, 2018), and *Hathewaya proteolytica* (UP000183952, July 9, 2018) UniProt databases simultaneously, using the HT-SEQUEST search algorithm, via the Proteome Discoverer (PD) Package (version 2.2, Thermo Scientific). Whole-tissue staged MS/MS data were also merged per donor as fractions and re-processed as above for whole-tissue/EV proteome correlation analyses. Ultracentrifugation libraries were included in PD processing of 15-fraction survey and EV-enriched pooled fraction experiments. Trypsin was set as the digestion enzyme while allowing up to four miss-cleavages, using 10 ppm precursor tolerance window and 0.6 Da fragment tolerance window. Oxidation of methionine and N-terminus acetylation were set as variable modifications, and carbamidomethylation of cysteine was set as a fixed modification. The peptide false discovery rate (FDR) was calculated using Percolator provided by PD and peptides were filtered based on a 1.0% FDR. Quantification utilized unique peptides (those assigned to a given Master protein group and not present in any other protein group) and razor peptides (peptides shared among multiple protein groups). Razor peptides were used to quantify only the protein with the most identified peptides and not for the other proteins they are contained in. A minimum of two unique peptides were required for a protein to be included in each dataset. To quantify peptide precursors detected in the MS1 but not sequenced from sample to sample, we enabled the ‘Feature Mapper’ node. Chromatographic alignment was done with a maximum retention time (RT) shift of 10 minutes and a mass tolerance of 10 ppm. Feature linking and mapping settings were: RT tolerance minimum of 0 minutes, mass tolerance of 10 ppm and signal-to-noise minimum of 5. Precursor peptide abundance quantification was based chromatographic intensities. For stand-alone 15-fraction survey experiments, there was no normalization of peptide amount/intensity. Total peptide amount was used for normalization in all other cases.

### RNA isolation

In EV-enriched pooled fraction experiments, the pellet resulting from ultracentrifugation of pooled fractions 1-4 was re-suspended in 700 μl of TRIzol (Invitrogen 15596026). Samples were then vortexed for 1 minute, and total RNA was extracted using the miRNeasy Mini Kit (Qiagen 217004) according to the manufacturer’s suggested protocol; each sample was eluted off the column twice using a total of 22 μl of RNase-free water.

### RNA sequencing

Fragment analysis was performed on a Bioanalyzer 2100 (Agilent) using the Small RNA Analysis Kit (Agilent DNF-470-0275). Resultant % miRNA composition is presented as mean ± standard error, and Student’s *t*-test (two-tailed, unpaired) was used for statistical comparison between tissue types with p ≤ 0.05 considered significant. cDNA libraries were synthesized with 4 ng of total RNA input to the Clontech SMARTer smRNA-seq kit (Takara Bio 635034) using the manufacturer’s suggested protocol. Small RNA libraries were size-purified using a Pippin Prep (Sage Science) selecting for library fragments ranging from 148-185bp on a 3% agarose gel. The finished libraries were quantified by Qubit fluorometer, Agilent TapeStation 2200, and RT-qPCR using the Kapa Biosystems library quantification kit according to manufacturer’s protocols. Uniquely indexed libraries were pooled in equimolar ratios targeting 20M clusters per library and two single-ended 75bp high-output (75 cycle) sequencing runs were performed on an Illumina NextSeq 500, to a target of 40 million reads per sample.

### Bioinformatic analyses

#### Proteomics

Bacterial proteins (originating from the *Clostridium*-derived collagenase) were excluded. For whole-tissue samples, collagenase libraries, and EV-enriched pooled fractions, the quantified proteins were exported from Proteome Discoverer and median-normalized (per sample) using in-house scripts written in Python v3.4.(21) Missing values were replaced with zero values in order to be analyzed by Qlucore Omics Explorer statistical software (Qlucore, v3.6): Thresholding of protein intensities was performed at level = 0.001, then intensities were log_2_-transformed for subsequent statistical analysis. In whole-tissue data, the ordinal disease stage was fit to a linear model of the stage factor as a predictor and retaining residuals. Significantly differentially-enriched proteins were calculated using a two-group comparison (ANOVA: independent measures, single factor design) at a Benjamini-Hochberg false discovery rate(48) (FDR, q or adjusted p-value) ≤0.05. PCA and heat map analyses (z-score-based, ordered by hierarchal clustering) were performed on the complete proteome (q ≤1). Per-protein normalized intensities from the overlapping portion of the merged whole-tissue and tissue-derived EV proteomes were averaged across donors, thresholded and transformed as above, then compared using linear regression and Pearson’s r correlation coefficient. Correlation significance was assessed by a two-tailed p-value at a 95% confidence interval in Prism (GraphPad, v5).

#### miRNA-Sequencing (miRNA-seq)

Data were retrieved from the sequencer in the form of concatenated FASTQ files, and processed using the open-source bcbio.nextgen framework (github.com/bcbio/bcbio-nextgen). In brief, the smallRNA-seq pipeline was utilized to detect adapter sequences via DNApi(49), which were removed by cutadapt.(50) STAR(51) was used to perform sequence alignment with the human genome, and seqcluster detected small RNA transcripts.(52, 53, 54) miRNAs were annotated using the miraligner tool with miRbase(55) as the reference miRNA database. Quality control was managed by FastQC.(56) After annotation, miRs were normalized, transformed, and tested for statistically-significant differential expression using DESeq2(57) defaults at an FDR ≤0.05. PCA and heat map analyses (z-score-based, ordered by q value) were performed on the complete miRNAome (q ≤1). We used TargetScan 7.2 to predict target genes for these EV miRs, with a high-confidence threshold of ≥ 95^th^ percentile weighted context++ score. After prediction of miR target genes, EV miRNA/mRNA target networks were generated in Gephi v0.9.2, where each identified miR was connected to its predicted targets.

#### Gene ontology, gene set/pathway enrichment and network analyses

Using ConsensusPathDB,(58) gene sets corresponding to the i) common whole-tissue proteome, ii) differentially-enriched whole-tissue proteins, iii) common EV proteome or miRNAome, iv) differentially-enriched EV proteins, and v) gene targets of differentially-expressed EV miRs were tested for enrichment by a hypergeometric test and adjusted for multiple comparisons using the Benjamini-Hochberg FDR. Pathways from BioCarta,(59) KEGG,(60) and Reactome(61) as well as gene ontology (GO) terms (retrieved from consensuspathdb.org in December 2019) with a p-value ≤0.05 were considered to be significantly-enriched in a gene set of interest. Significant overrepresentation of vesicle-associated pathways in the common whole-tissue proteome was assessed by comparing incidence of pathway names that included “vesicle” in the enriched pathways list (7/484) vs. the total pathway database (7/2,459) using a two-tailed Chi-square test at p ≤0.05. GO term enrichment of the common whole-tissue proteome was assessed using terms found in the biological process, molecular function, and cellular component categories for all GO levels. Coabundance profiling in 15-fraction survey experiments was performed using high-dimensional quantitative clustering in XINA (v3.9).(22, 46) Raw protein intensities from PD2.2 were normalized by total protein intensity per donor. Normalized protein abundances per OptiPrep fraction were merged into two tissue-specific datasets, each with three fraction ranges (F1-4, 5-10, 11-15) for subsequent clustering analyses. Coabundance patterns were fit to 10 clusters per analysis and functionality of cluster constituents was assessed by ConsensusPathDB as above.

Pathway networks are composed of pathways as the nodes and shared genes between pathways as the edges. Node size corresponds to –log (q-value) and edge weight (thickness) corresponds to the gene overlap between pairs of pathways measured by the Jaccard index J, which is defined as

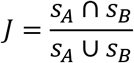

where *s*_*A*_ and *s*_*B*_ are the set of proteins detected in proteomics and gene targets of miRs detected in transcriptomics that belong to pathway *A* and pathway *B*, respectively. Edges with a Jaccard index < 0.1 were discarded in the visualization for clarity. Modularity optimization via the Louvain method(25) was utilized to cluster pathway nodes into real network communities, overarching community functions were manually assigned, and standalone nodes were removed. The network visualizations were made using Gephi v0.9.2. Fisher’s exact test (two-tailed) was used to assess overlap of pathways that were significantly enriched in both the common EV proteome and amongst gene targets of the common EV miRNAome, which was considered to be significant at p ≤ 0.05. Integrated pathway-specific PPI networks were produced by mapping the gene list that defined the pathway of interest from ConsensusPathDB onto the STRING v11.0 (62) PPI network (at a confidence score cutoff of 800). Nodes (and their shared edges) that were significantly enriched between tissue types in the EV proteome or that were high-confidence targets of miRs significantly enriched between tissue types in the EV miRNAome were highlighted. STRING v11.0 was also used at default settings with disconnected nodes hidden to generate protein-protein interaction networks, GO cellular component, UniProt Keyword, and KEGG pathway analyses of the impacts of OptiPrep density gradient separation and targeted mass exclusion on extracellular vesicle purification.

## Declarations

### Ethics Approval and Consent to Participate

All human samples were handled in accordance with Institutional Review Board (IRB) protocols to S.C.B, J.D.M. (2011P001703), and Peter Libby (P.L.; 1999P001348), approved by the Partners Human Research Committee/Brigham and Women’s Hospital IRB. Sample collection included written consent for publication and use of samples from all donors. All experimental methods used comply with the Declaration of Helsinki.

## Consent for Publication

Not applicable.

## Author Contributions

M.C.B, F.B., and E.A. conceptualized the study. M.C.B, F.B., A.H., F.S., M.A.R., and S.A.S. conceived and designed the experiments. M.C.B., F.B., A.H., F.S., H.H., L.P.R., L.A.S., S.K.A., T.P., and S.A.S. contributed to the collection and analysis of data. E.S., G.K.S., S.C.B., and J.D.M. provided tissue. M.C.B. wrote the manuscript. S.M., G.C., M.A., and E.A. provided financial support and critically reviewed the manuscript. All authors contributed to revising and editing of the manuscript.

## Acknowledgments

We gratefully acknowledge Dr. Peter Libby (Center for Excellence in Vascular Biology, Brigham and Women’s Hospital, Harvard Medical School) for provision of carotid endarterectomy tissue. We thank Maria Ericsson at the Harvard Medical School Electron Microscopy Facility for assistance with electron microscopy, and Svetlana Gorbatov, James Gosnell, and Amanda Pang for assistance in collecting clinical samples. Sequencing was performed by the Molecular Biology Core Facilities at Dana-Farber Cancer Institute, and we thank Dr. Gary Sommerville and Dr. Zachary Herbert for their assistance.

