## Supplemental Figures 1-14 for "Conserved and Divergent Modulation of Calcification in Atherosclerosis and Aortic Valve Disease by Tissue Extracellular Vesicles"

**Supplemental Figure 1: OptiPrep Density Gradient Separation to Enrich Cardiovascular Tissue-Entrapped EVs. A,** Human carotid artery plaques or calcified aortic valves from patients diagnosed with calcific aortic valve disease (CAVD) were surgically removed and promptly underwent rough dissection and enzymatic digestion with bacterial collagenase to dissociate the tissue components. Low-speed ultracentrifugation collected cells and debris, and the supernatant underwent high-speed centrifugation to pellet large microsomes and apoptotic bodies. The resultant supernatant was ultracentrifuged twice, with a wash step in between. The ultracentrifuged pellet (rich in EVs, as well as ECM, ribosomal debris, and protein aggregates) was overlaid on a linear 10-30% isosmotic iodixanol gradient, and OptiPrep density gradient separation was performed. **B,** In 15-fraction survey experiments, each OptiPrep fraction was collected and pelleted separately by ultracentrifugation. Every fraction then underwent mass spectrometry, nanoparticle tracking analysis, and transmission electron microscopy. **C,** In EV-enriched OptiPrep fraction experiments, fractions 1-4 were pooled and pelleted together. High-resolution mass spectrometry, miRNA-seq, nanoparticle tracking analysis, electron microscopy, and alkaline phosphatase activity assays were completed on the pooled EV-enriched fractions.

**Supplemental Figure 2: EV Enrichment by OptiPrep Separation – EV Markers and XINA Co-abundance Profiling. A,** From 15-fraction survey experiments on human carotid artery plaques (left) and calcified aortic valves (right), protein abundance heat maps of 22 EV markers across 15 OptiPrep density gradient fractions found that EV marker proteins were consistently enriched in the four least-dense fractions of both tissue types; n=3 carotid and 4 aortic valve donors. **B,** Tissue-specific XINA coabundance profiling of protein abundances in OptiPrep fractions 1-4 from carotid artery plaques (left) and calcified aortic valves (right). Proteins detected in the 5 clusters per tissue type whose protein abundance profiles mimicked those of the 22 EV markers in part A (elevated in fractions 2 and 3; brown clusters 1, 2, 5, 6, 9 and 1, 5, 6, 8, 10 for carotid artery and aortic valve, respectively) were subjected to gene ontology analysis. Highly significant GO terms in these clusters were largely associated with vesicular processes and further demonstrated EV enrichment in the least-dense OptiPrep fractions.

**Supplemental Figure 3: Transmission Electron Microscopy of 15-Fraction Survey Experiments**. **A and B,** Representative CD63-labelled immunogold transmission electron microscopy (TEM) and negative controls from OptiPrep fractions 1-15 of human carotid artery (A) and calcified aortic valve (B) demonstrated fraction density-dependent enrichment of extracellular vesicles, globular collagen, and fibrillar collagen.

**Supplemental Figure 4: Transmission Electron Microscopy Negative Controls.** Representative negative control transmission electron microscopy (corresponding to Figure 3C) identified membrane-bound EVs in OptiPrep fractions 1-4 (arrows) from carotid artery plaques (top) and calcified aortic valve (bottom); bar=100 nm. Consistent with mass spectrometry-derived protein abundance, TEM showed abundant globular collagens in fractions 5-10 (arrowheads) and fibrillar collagens in the most-dense fractions (open arrows).

**Supplemental Figure 5: Tissue-Entrapped EVs Purified by Ultracentrifugation Alone Show Marked Extracellular Matrix Contamination. A,** Purification of extracellular vesicles (EVs) was assessed by comparing the proteome collected by ultracentrifugation alone vs. EVs enriched by pooling fractions 1-4 of an OptiPrep density gradient. **B,** Proteomics found that the OptiPrep-derived EV proteome was >38% deeper than that of EVs obtained by ultracentrifugation alone in both carotid artery (n=4; 39.7% larger) and aortic valve tissues (n=4; 38.1% larger). **C,** Top 10 gene ontology (cellular component) and UniProt keyword analyses of those proteins unique to ultracentrifugation found an abundance of extracellular matrix and endoplasmic reticulum protein contaminants when tissue-derived EVs were extracted by ultracentrifugation alone. Meanwhile, OptiPrep enrichment enabled detection of additional intracellular-, membrane- and vesicle-associated proteins linked with diverse biological functions (e.g. phosphorylation, protein transport, post-translational modification, actin-binding, etc.).

**Supplemental Figure 6: Elimination of Iodixanol From Mass Spectrometry by Targeted Mass Exclusion Windows. A,** Total ion current spiked during mass spectrometry of samples prepared by OptiPrep density gradient (gold shading: representative spiking at 28-34 minutes during a 90-minute gradient). **B,** MS scans from 28-34 minutes of a 90-minute gradient, showing a highly-abundant precursor ion (775.86354, z=2, purple shading) consistent with carryover of iodixanol contamination from the OptiPrep gradient (theoretical mass=1549.7133 u). **C,** The 775.8 m/(z precursor eluted repeatedly between 28-34 minutes; inset: peaks in the range of ~775.8 m/z were limited to this RT window. **D,** Despite dynamic exclusion of identical precursor ions, peptide sequencing events were frequently performed on fragment ions from the 775.86354 m/z iodixanol contaminant. **E,** Flow chart describing the peptide sequencing approach that was subsequently employed to avoid wasting sequencing events on iodixanol precursor ions. Targeted mass exclusion was enacted at an m/z of 775.8645 (z=2), an exclusion mass width of 10 ppm, over a 4 minute (30-minute gradients) or 7-minute (90-minute gradients) retention time window. **F,** The total number of proteins detected increased by 9.5% with targeted mass exclusion of EV-enriched pooled fractions from 4 carotid artery and 4 aortic valve donors (955 vs. 1,046 proteins). **G,** The 198 proteins unique to targeted mass exclusion comprised a tightly associated protein-protein interaction network and 20 significantly enriched KEGG pathways, while those lost during targeted exclusion had no significant pathway enrichment.

**Supplemental Figure 7: Mock Iodixanol-Only OptiPrep Fractions. A,** Mock OptiPrep fractions prepared in the absence of ultracentrifuged tissue digest (i.e. containing only iodixanol and NTE buffer) continued to demonstrate spikes in full MS total ion current (gold shading: representative spiking at 17-21 minutes during a 30-minute gradient). **B,** The 775.86343 m/z precursor ion (purple shading) continued to be highly abundant during this retention time window. **C,** Peaks in the range of ~775 m/z were limited to this time period. Together, this data indicated that these peaks were not biological peptides derived from cardiovascular tissue, but rather residual iodixanol contamination.

**Supplemental Figure 8: Labelled Integrated Network of Shared Proteomic and Transcriptomic Pathways Enriched in Tissue-Entrapped EVs.** A statistically significant number of KEGG, Reactome, and BioCarta pathways (154 pathways, p=0.000206) were significantly enriched in both the proteome and gene targets of the miRNAome that was common to carotid artery and aortic valve-derived EVs (n=4). The network of these overlapping pathways was generated with pathways as the nodes (node size corresponds to -log(q-value)) and shared detected genes between pathways as the edges (edge thickness matches the Jaccard index of overlap between detected genes of the two connected pathway nodes). Unbiased clustering of pathways into real network communities by the Louvain method revealed 9 distinct functions shared by cardiovascular tissue-derived EV cargoes, including cell cycle regulation, synthesis and organization of the extracellular matrix (ECM), and modulation of MAPK and Rho GTPase intracellular signaling cascades.

**Supplemental Figure 9: “Focal Adhesion” Pathway-Specific Protein-Protein Interaction Network from Integrated Tissue EV Multi-Omics.** Overlap of pathways significantly associated with proteins and gene targets of miRs that were differentially enriched between carotid artery and aortic valve-derived EVs was assessed (n=4). Pathway-specific protein-

protein interaction networks were utilized to integrate EV multi-omics. The KEGG “Focal Adhesion” pathway (hsa04510) was significantly enriched in multiple layers of carotid artery-derived EV omics. Node diameter corresponds to node degree.

**Supplemental Figure 10: “Regulation of Actin Cytoskeleton” Pathway-Specific Protein-Protein Interaction Network from Integrated Tissue EV Multi-Omics.** Overlap of pathways significantly associated with proteins and gene targets of miRs that were differentially enriched between carotid artery and aortic valve-derived EVs was assessed (n=4). Pathway-specific protein-protein interaction networks were utilized to integrate EV multi-omics. The KEGG “Regulation of Actin Cytoskeleton” pathway (path:hsa04810) was significantly enriched in multiple layers of carotid artery-derived EV omics. Node diameter corresponds to node degree.

**Supplemental Figure 11: “Hypertrophic Cardiomyopathy” Pathway-Specific Protein-Protein Interaction Network from Integrated Tissue EV Multi-Omics.** Overlap of pathways significantly associated with proteins and gene targets of miRs that were differentially enriched between carotid artery and aortic valve-derived EVs was assessed (n=4). Pathway-specific protein-protein interaction networks were utilized to integrate EV multi-omics. The KEGG “Hypertrophic Cardiomyopathy” pathway (path:hsa05410) was significantly enriched in multiple layers of carotid artery-derived EV omics. Node diameter corresponds to node degree.

**Supplemental Figure 12: “Dilated Cardiomyopathy” Pathway-Specific Protein-Protein Interaction Network from Integrated Tissue EV Multi-Omics.** Overlap of pathways significantly associated with proteins and gene targets of miRs that were differentially enriched between carotid artery and aortic valve-derived EVs was assessed (n=4). Pathway-specific protein-protein interaction networks were utilized to integrate EV multi-omics. The KEGG

“Dilated Cardiomyopathy” pathway (path:hsa05414) was significantly enriched in multiple layers of carotid artery-derived EV omics. Node diameter corresponds to node degree.

**Supplemental Figure 13: “Viral Myocarditis” Pathway-Specific Protein-Protein Interaction Network from Integrated Tissue EV Multi-Omics.** Overlap of pathways significantly associated with proteins and gene targets of miRs that were differentially enriched between carotid artery and aortic valve-derived EVs was assessed (n=4). Pathway-specific protein-protein interaction networks were utilized to integrate EV multi-omics. The KEGG “Viral Myocarditis” pathway (path:hsa05416) was significantly enriched in multiple layers of carotid artery-derived EV omics. Node diameter corresponds to node degree.

**Supplemental Figure 14: “Amino Sugar and Nucleotide Sugar Metabolism” Pathway-Specific Protein-Protein Interaction Network from Integrated Tissue EV Multi-Omics.** Overlap of pathways significantly associated with proteins and gene targets of miRs that were differentially enriched between carotid artery and aortic valve-derived EVs was assessed (n=4). Pathway-specific protein-protein interaction networks were utilized to integrate EV multi-omics. The KEGG “Amino Sugar and Nucleotide Sugar Metabolism” pathway (path:hsa00520) was significantly enriched in multiple layers of carotid artery-derived EV omics. Node diameter corresponds to node degree.

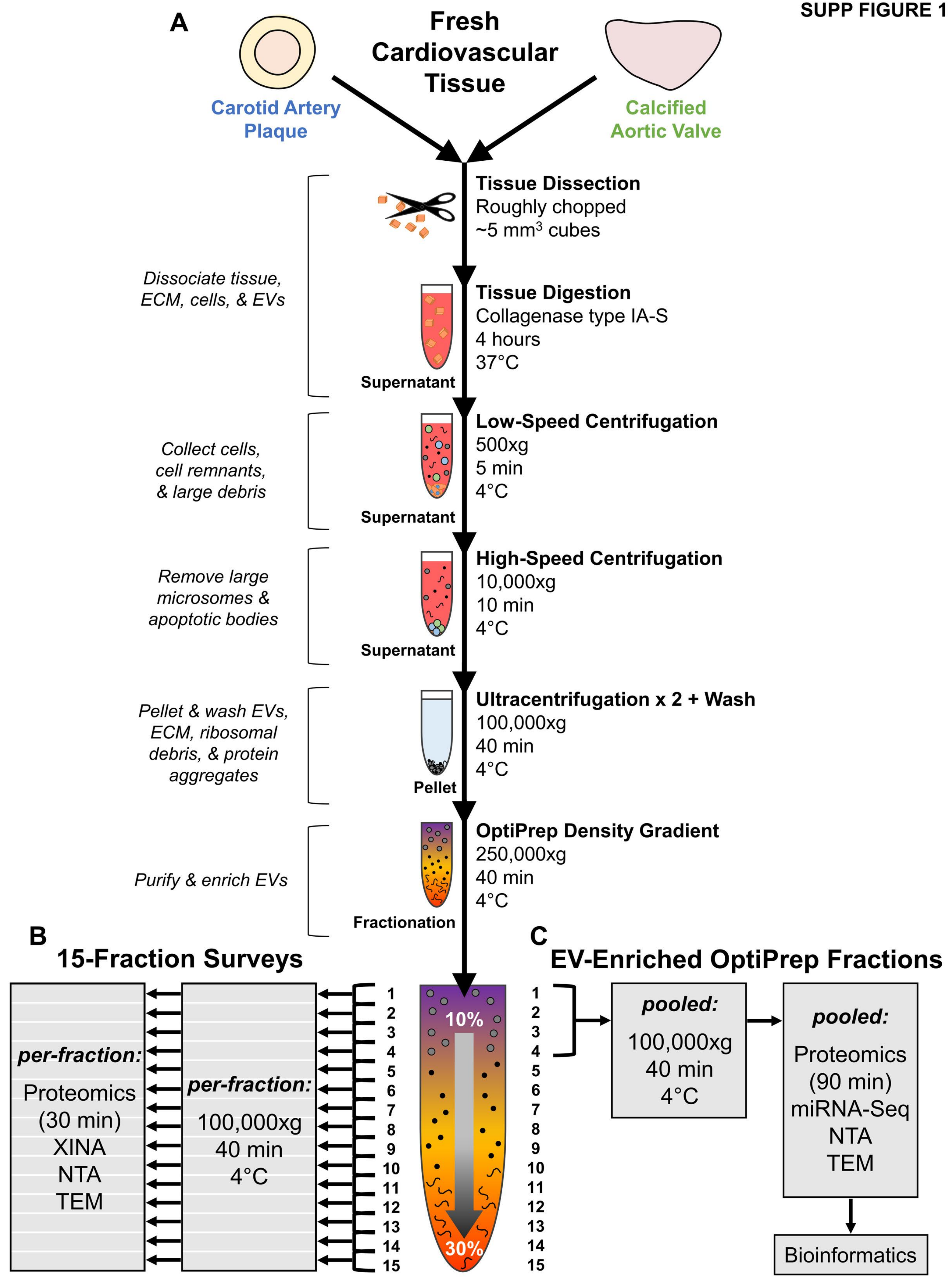
A
Fresh Cardiovascular Tissue
Carotid Artery Plaque
Calcified Aortic Valve
Tissue Dissection
Roughly chopped
~5 mm3 cubes
Dissociate tissue, ECM, cells, & EVs
Tissue Digestion
Collagenase type IA-S
4 hours
37°C
Supernatant
Collect cells, cell remnants, & large debris
Low-Speed Centrifugation
500xg
5 min
4°C
Supernatant
Remove large microsomes & apoptotic bodies
High-Speed Centrifugation
10,000xg
10 min
4°C
Supernatant
Pellet & wash EVs, ECM, ribosomal debris, & protein aggregates
Ultracentrifugation x 2 + Wash
100,000xg
40 min
4°C
Pellet
Purify & enrich EVs
OptiPrep Density Gradient
250,000xg
40 min
4°C
Fractionation
B
15-Fraction Surveys
per-fraction:
Proteomics
(30 min)
XINA
NTA
TEM
per-fraction:
100,000xg
40 min
4°C
1 2 3 4 5 6 7 8 9 10 11 12 13 14 15
10%
30%
C
EV-Enriched OptiPrep Fractions
pooled:
100,000xg
40 min
4°C
pooled:
Proteomics
(90 min)
miRNA-Seq
NTA
TEM
Bioinformatics

SUPP FIGURE 2

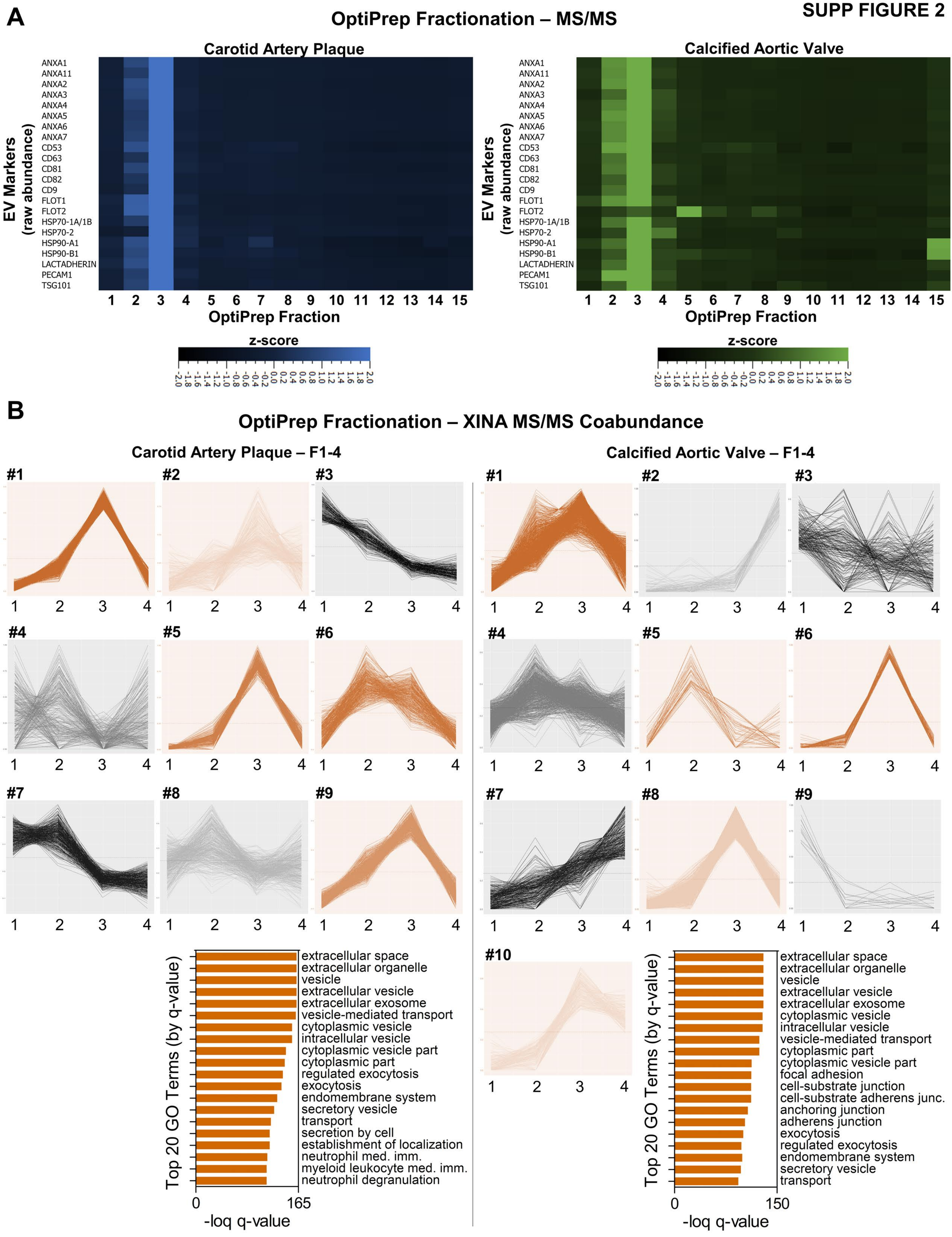

SUPP FIGURE 3

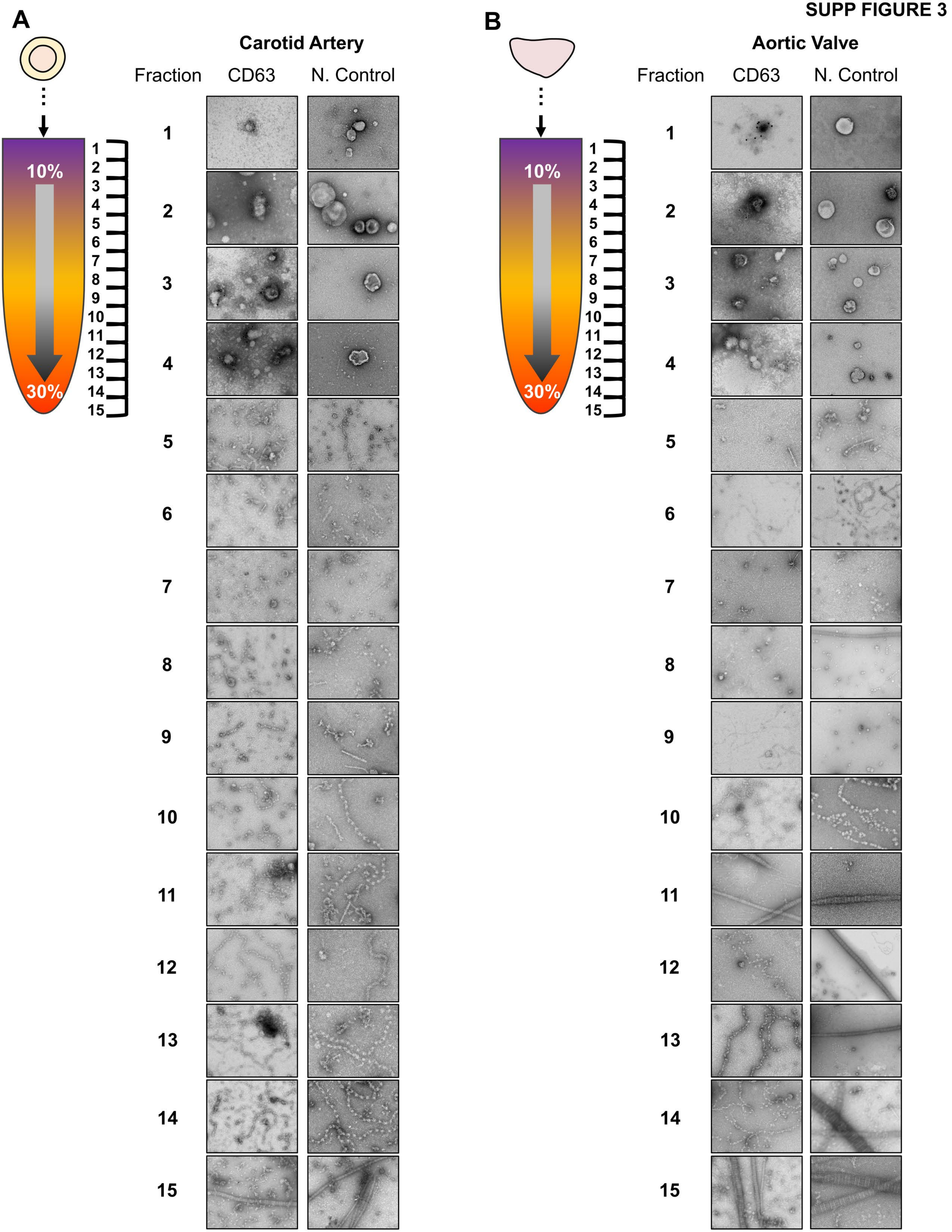

SUPP FIGURE 4

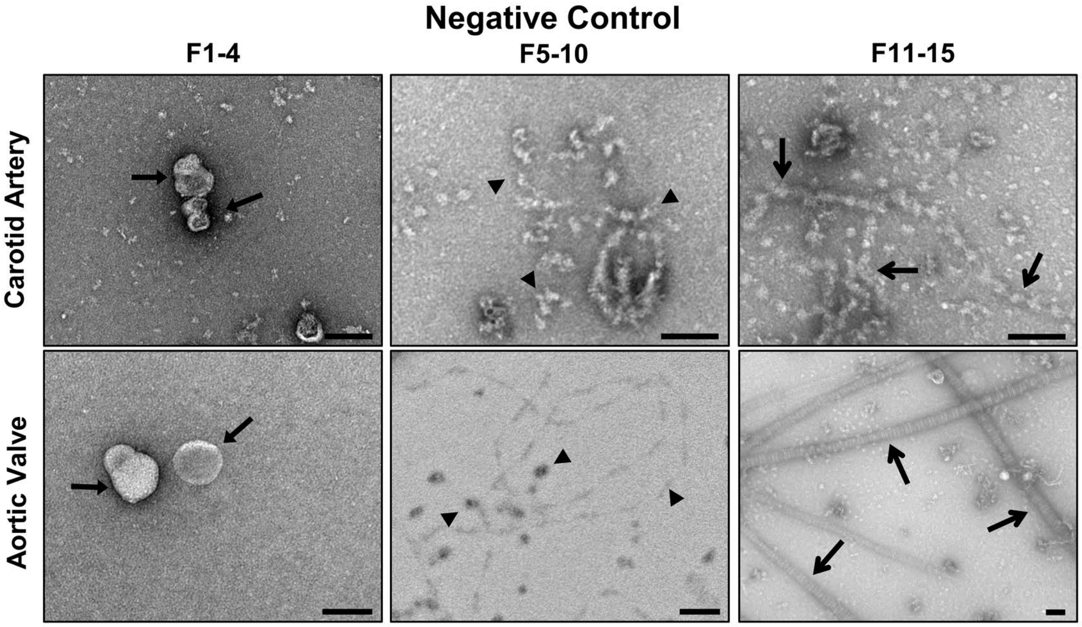

SUPP FIGURE 5

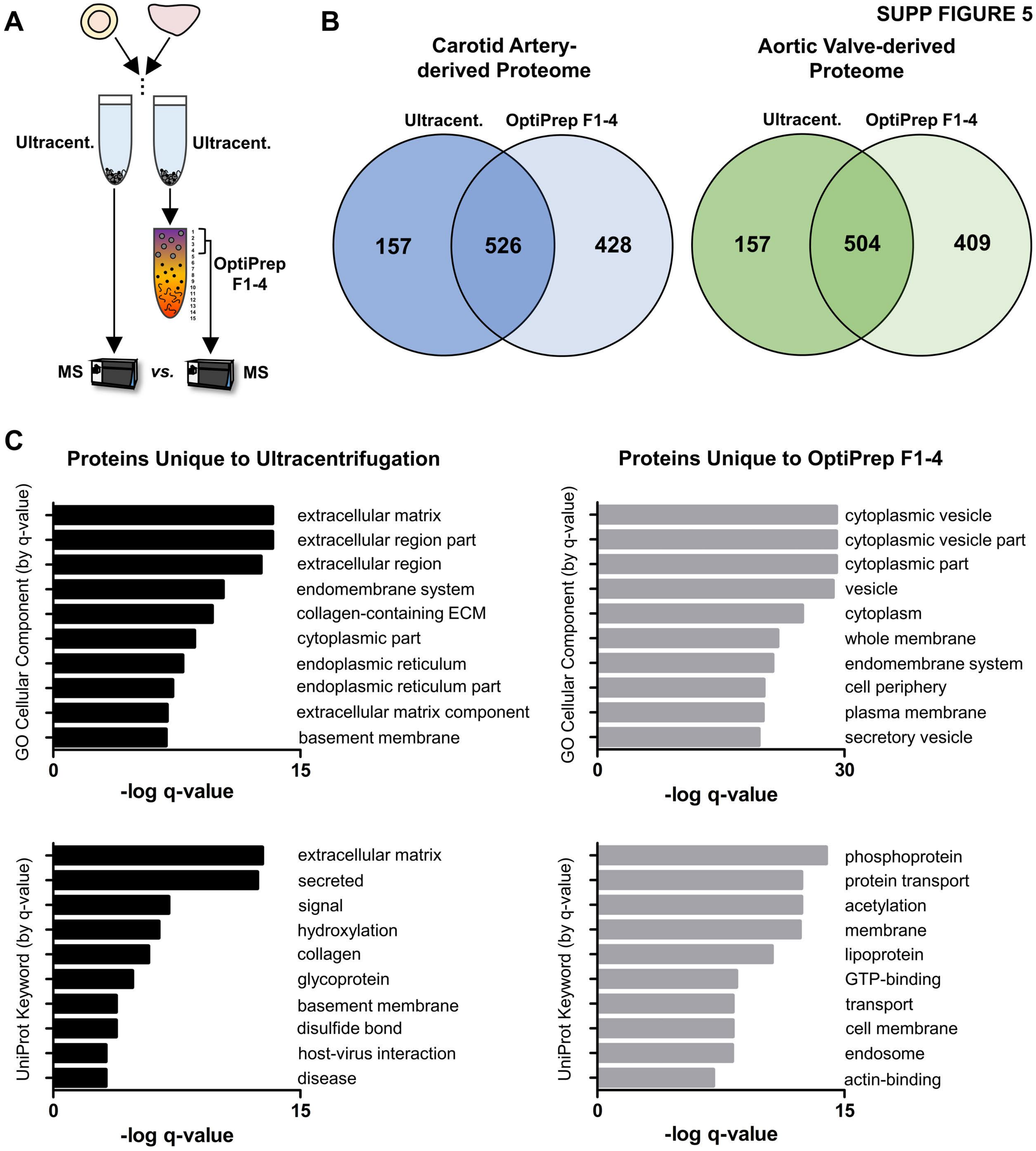

**SUPP FIGURE 6**

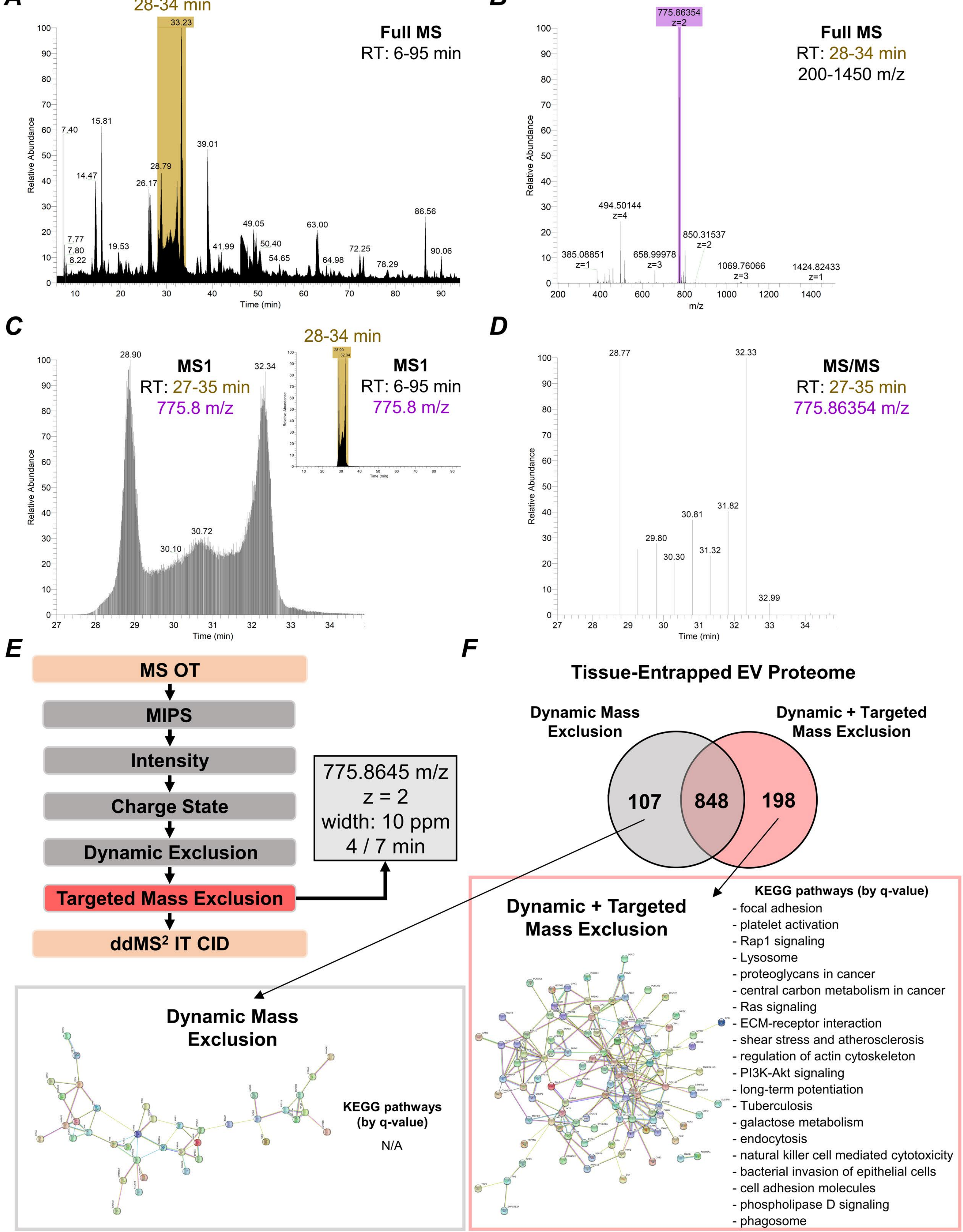

A

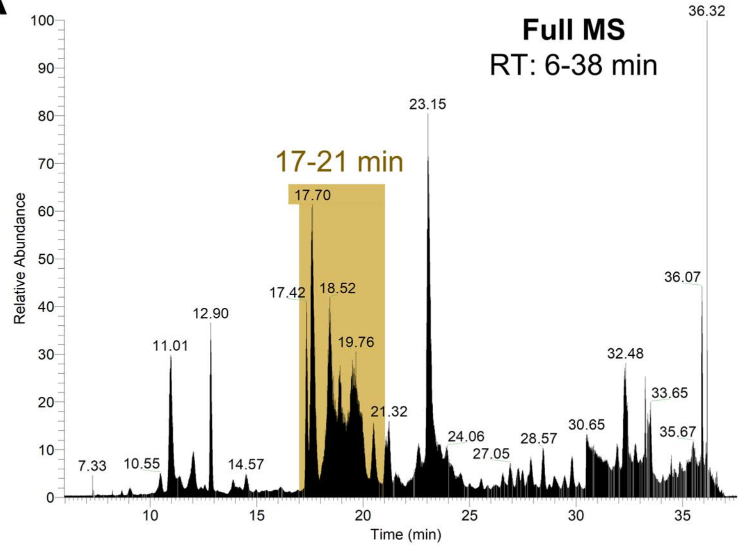
Full MS
RT: 6-38 min
17-21 min
Relative Abundance
Time (min)
36.32
23.15
17.70
17.42
18.52
19.76
12.90
11.01
21.32
36.07
32.48
33.65
35.67
30.65
28.57
27.05
24.06
14.57
10.55
7.33

B

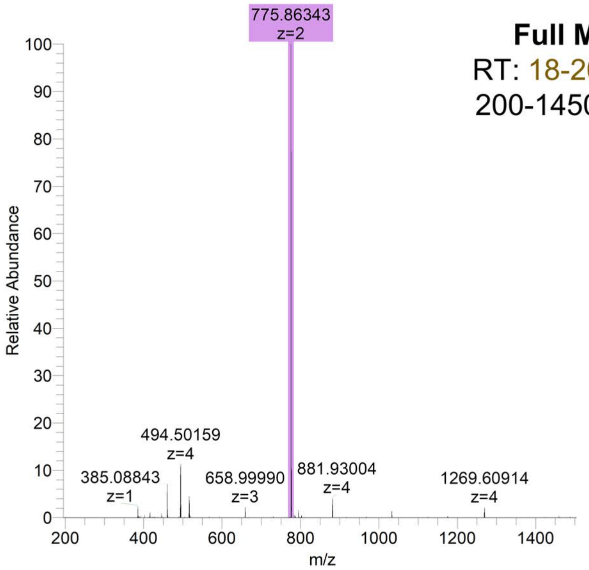
Full MS
RT: 18-20 min
200-1450 m/z
775.86343
z=2
494.50159
z=4
385.08843
z=1
658.99990
z=3
881.93004
z=4
1269.60914
z=4
Relative Abundance
m/z

C

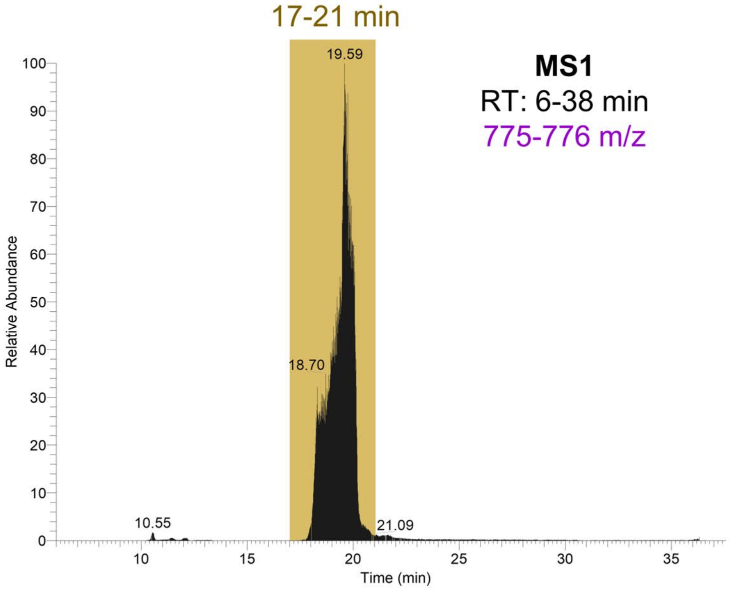
17-21 min
MS1
RT: 6-38 min
775-776 m/z
19.59
18.70
10.55
21.09
Relative Abundance
Time (min)

### SUPP FIGURE 8

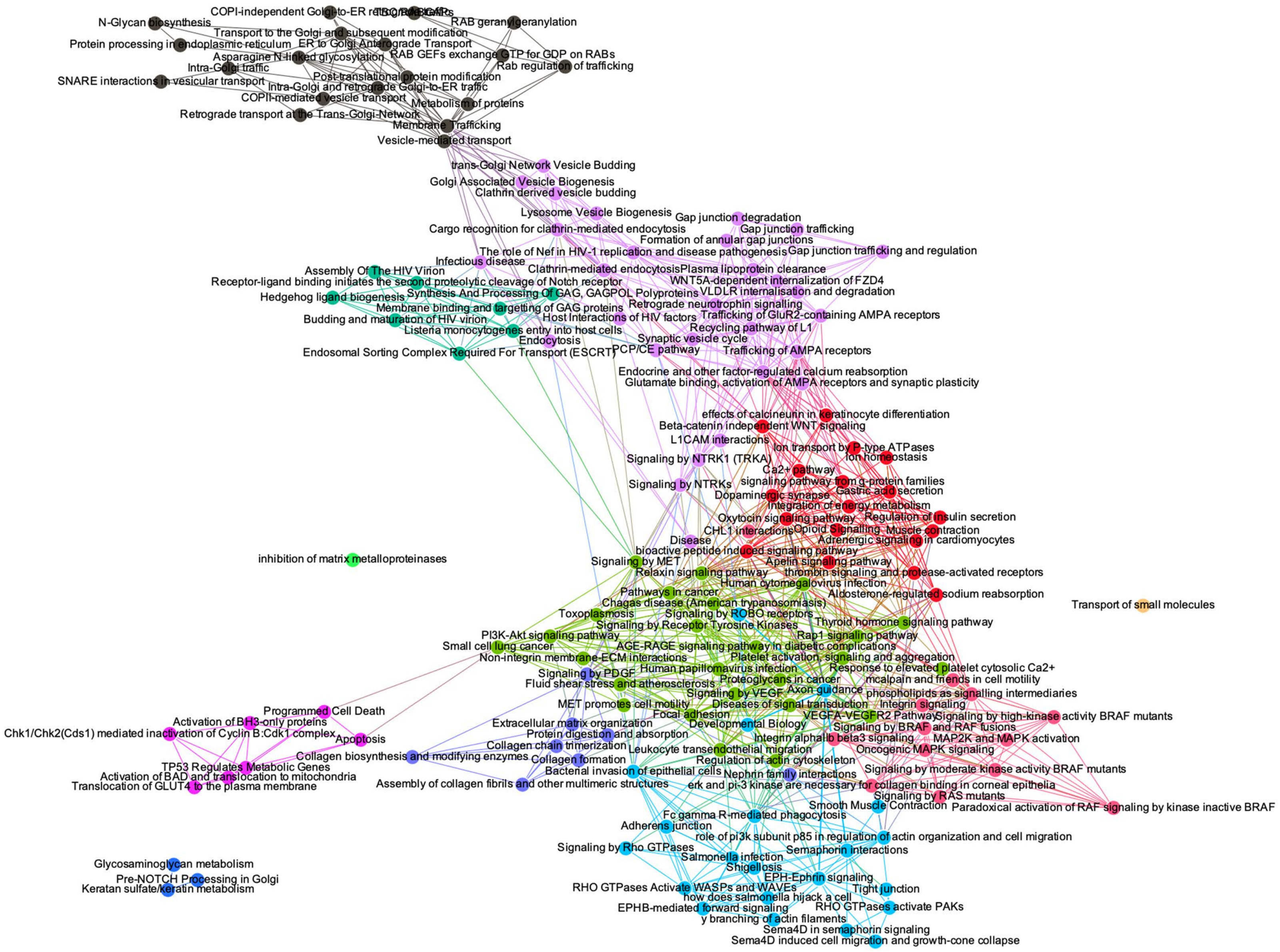

#### SUPP FIGURE 9

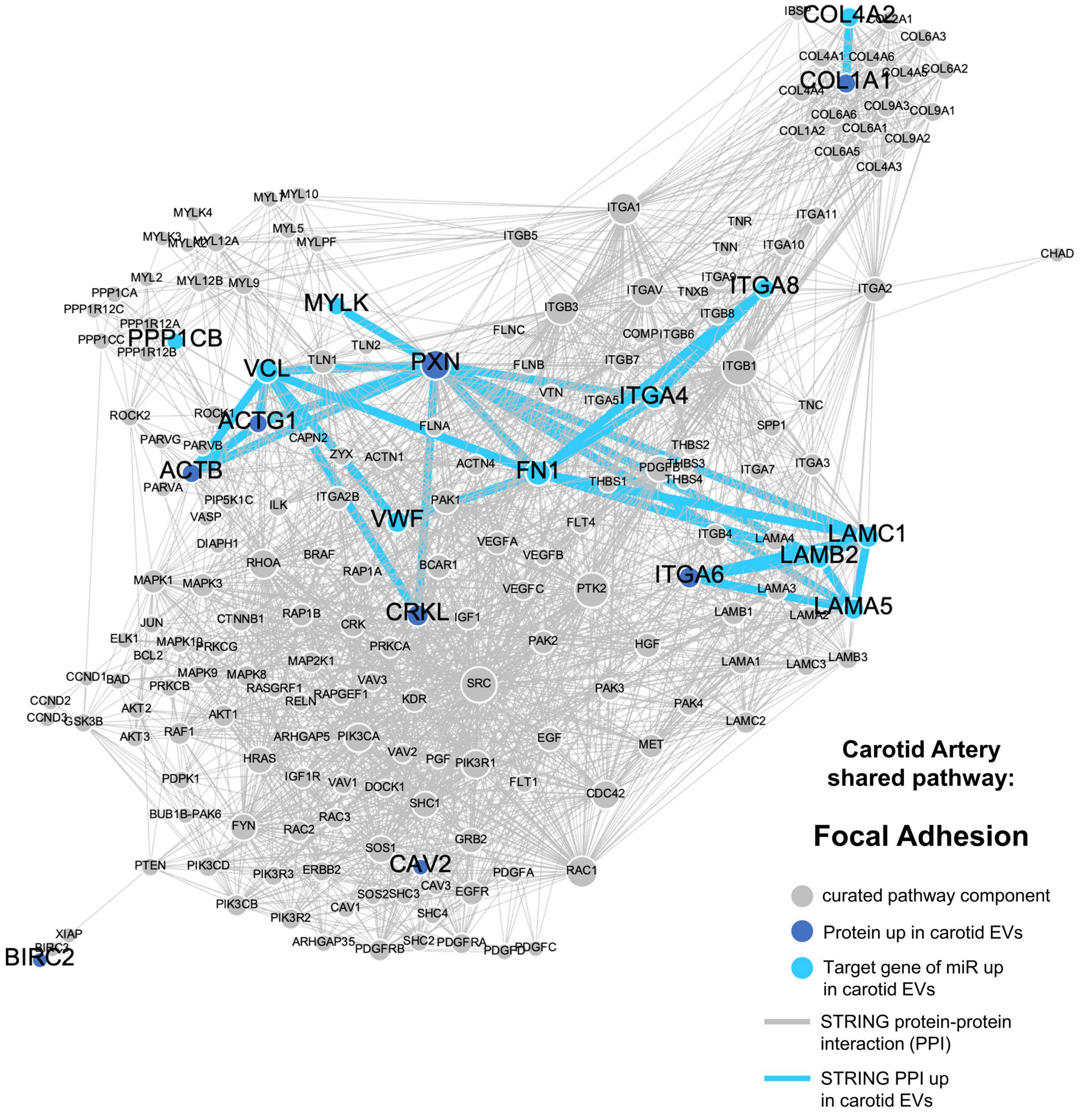

**SUPP FIGURE 10**

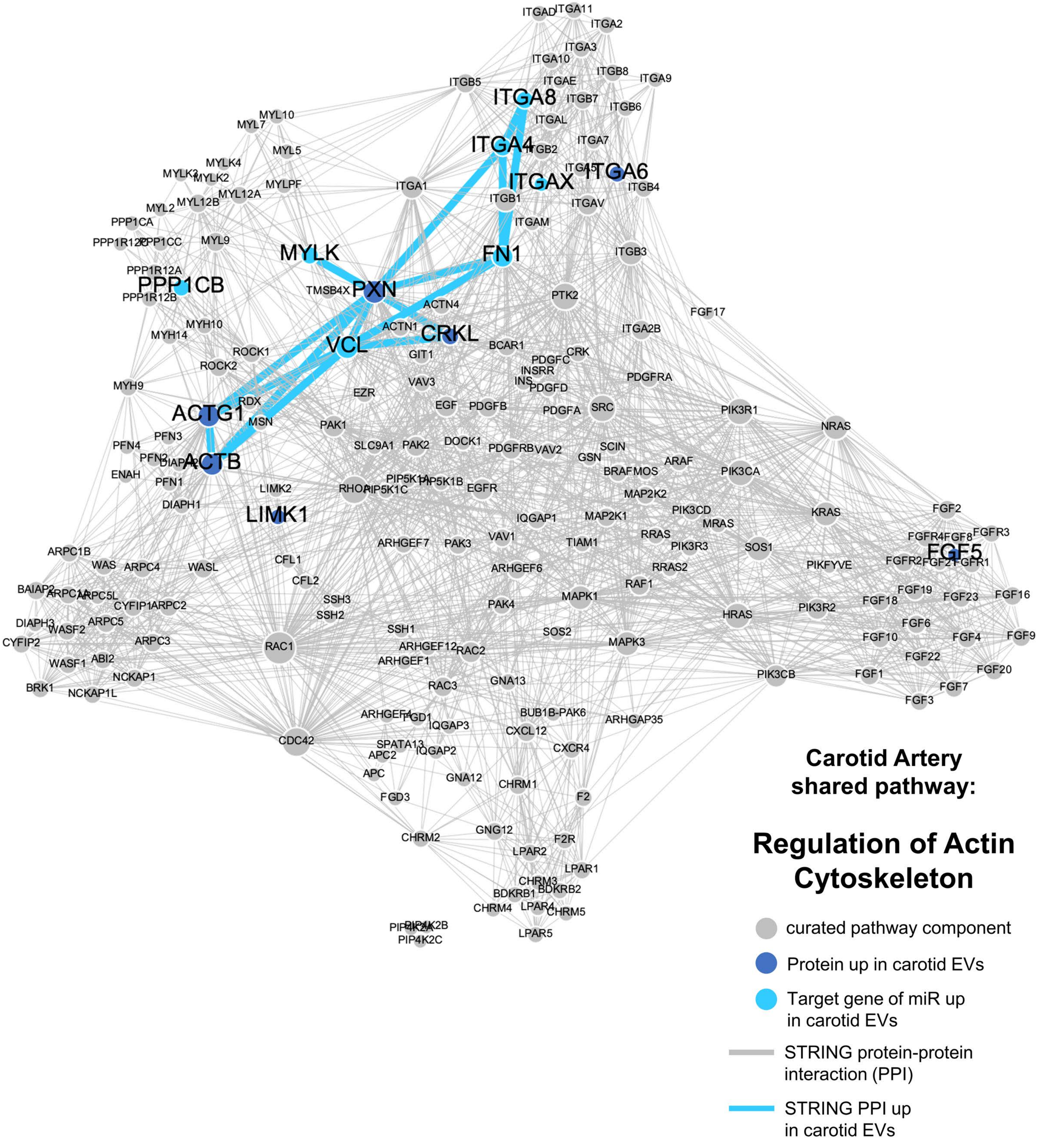

**SUPP FIGURE 11**

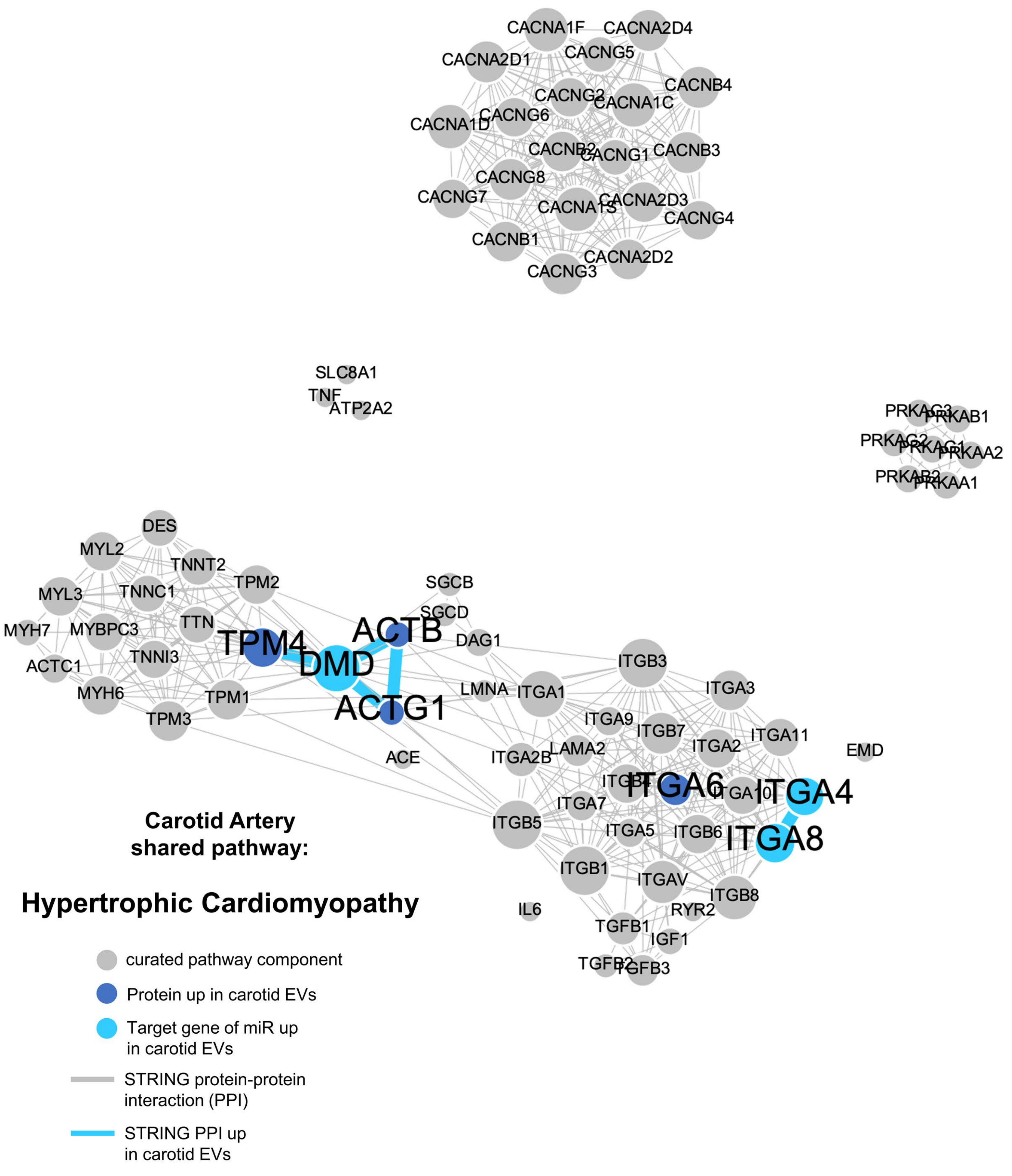

**SUPP FIGURE 12**

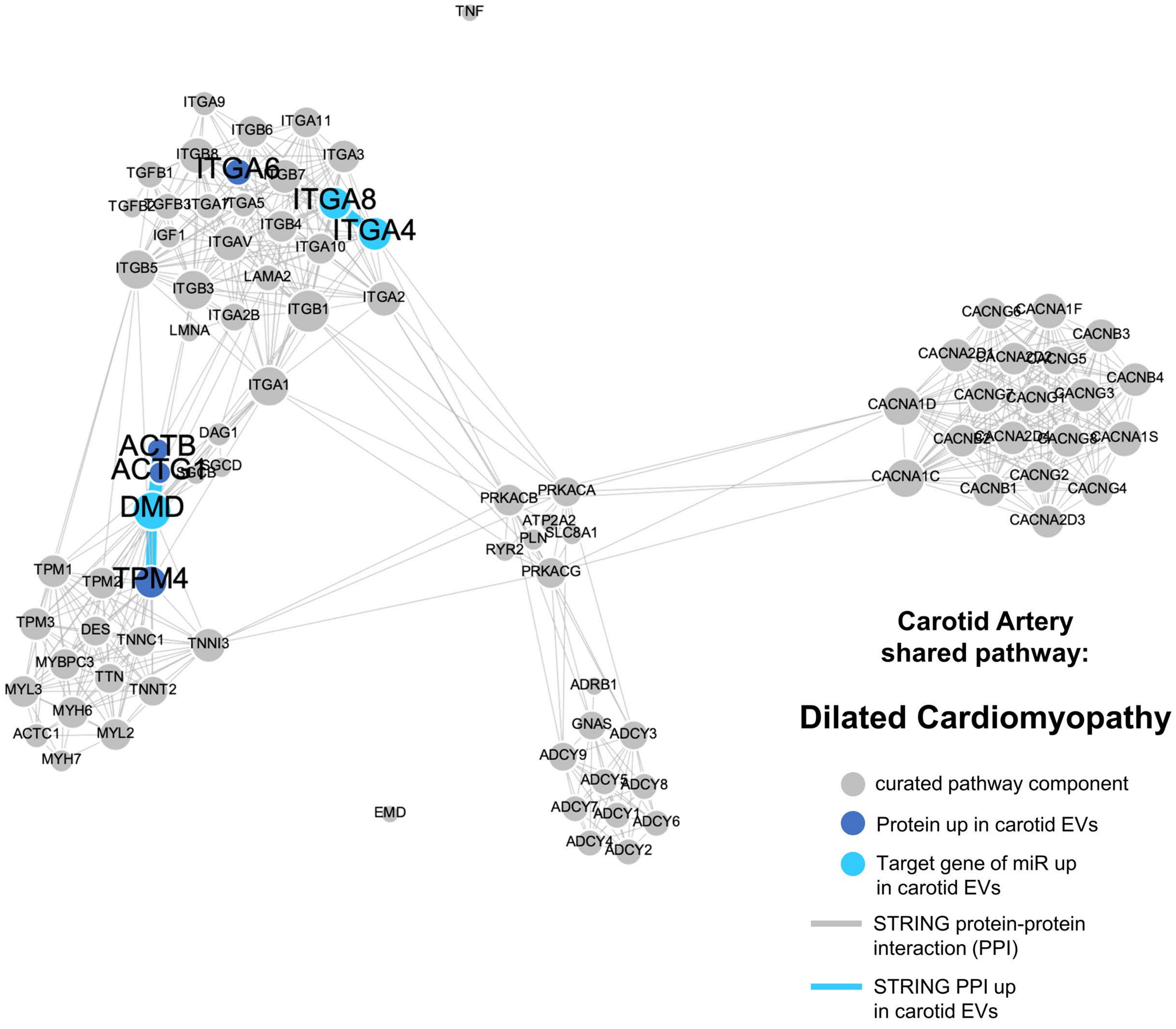

**SUPP FIGURE 13**

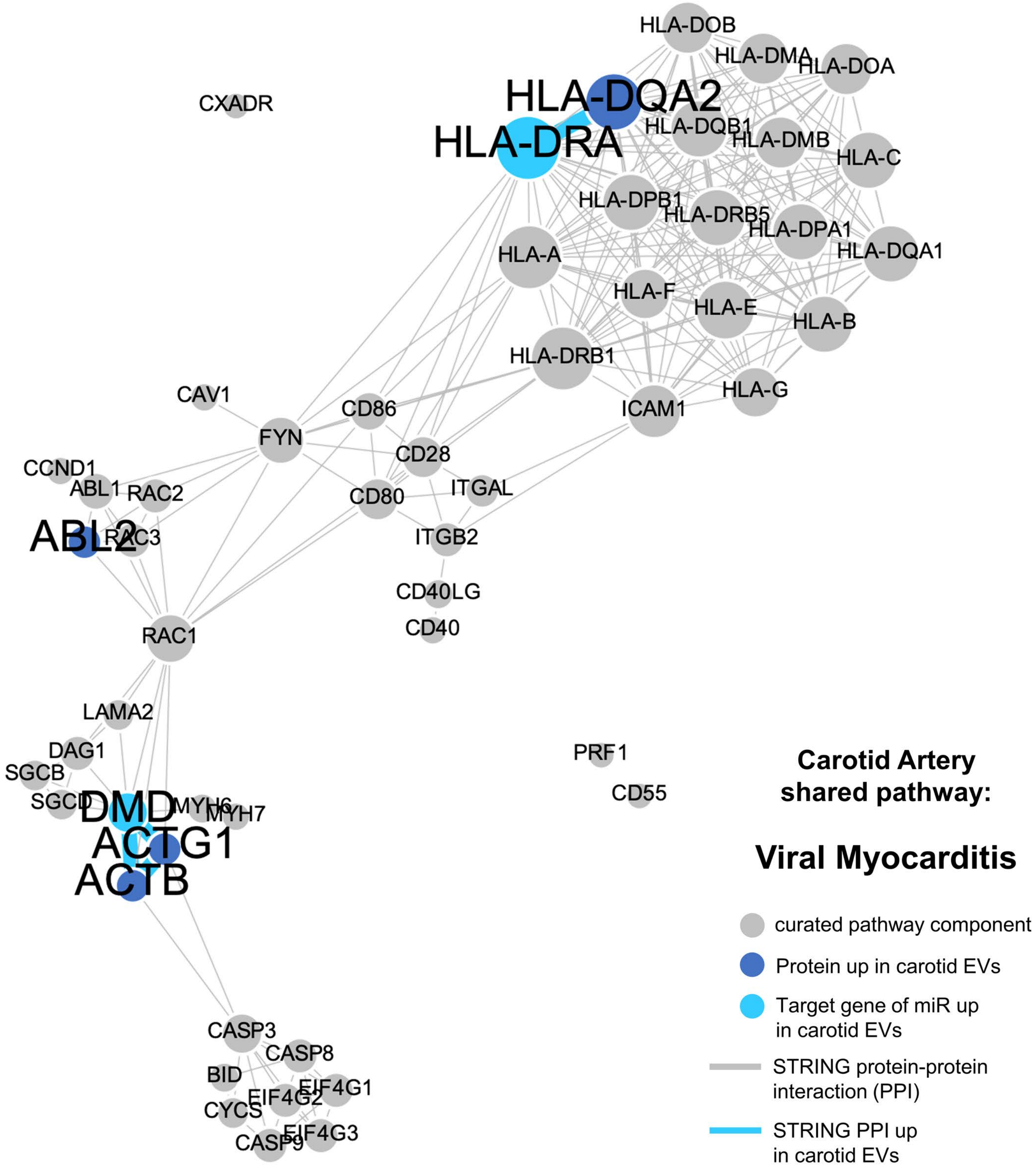

**SUPP FIGURE 14**

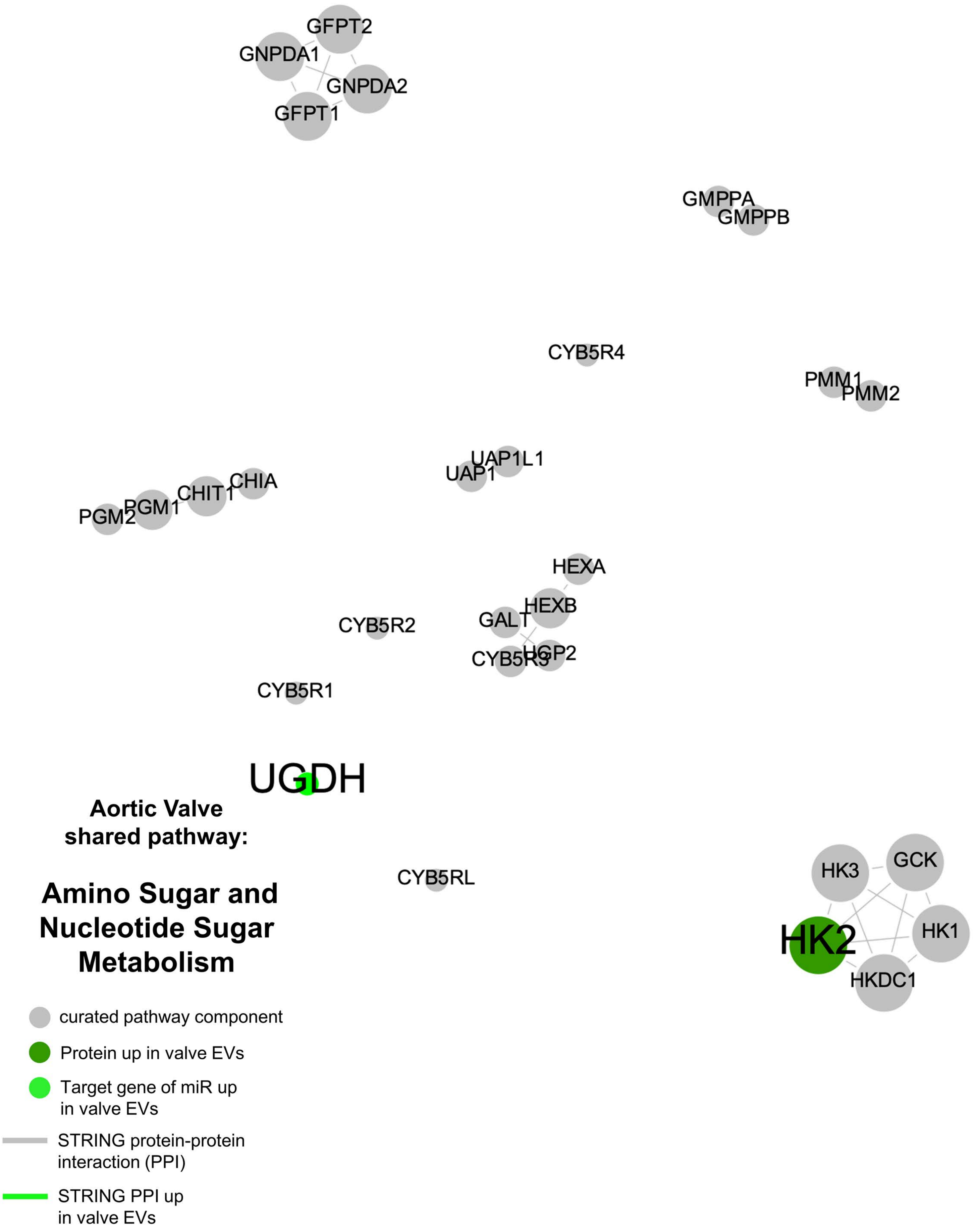
